# *In vitro* production of cat-restricted *Toxoplasma* pre-sexual stages by epigenetic reprogramming

**DOI:** 10.1101/2023.01.16.524187

**Authors:** Ana Vera Antunes, Martina Shahinas, Christopher Swale, Dayana C. Farhat, Chandra Ramakrishnan, Christophe Bruley, Dominique Cannella, Charlotte Corrao, Yohann Couté, Adrian B. Hehl, Alexandre Bougdour, Isabelle Coppens, Mohamed-Ali Hakimi

## Abstract

Sexual reproduction of *Toxoplasma gondii*, which is restricted to the small intestine of felids, is sparsely documented, due to ethical concerns surrounding the use of cats as model organisms. Chromatin modifiers dictate the developmental fate of the parasite during its multistage life cycle, but their targeting to stage-specific cistromes is poorly described^1^. In this study, we found that transcription factors AP2XII-1 and AP2XI-2, expressed in tachyzoite stage that causes acute toxoplasmosis, can silence genes necessary for merozoites, a developmental stage critical for sexual commitment and transmission to the next host, including humans. Their conditional and simultaneous depletion leads to a drastic change in the transcriptional program, promoting a complete transition from tachyzoites to merozoites. Pre-gametes produced *in vitro* under these conditions are characterized by specific protein markers and undergo typical asexual endopolygenic division cycles. In tachyzoites, AP2XII-1 and AP2XI-2 bind DNA as heterodimers at merozoite promoters and recruit the epigenitors MORC and HDAC3^1^, which in turn restrict the accessibility of chromatin to the transcriptional machinery. Thus, the commitment to merogony stems from a profound epigenetic rewiring orchestrated by AP2XII-1 and AP2XI-2. This effective *in vitro* culture of merozoites paves the way to explore *Toxoplasma* sexual reproduction without the need to infect kittens and has potential for the development of therapeutics to block parasite transmission.

---

The parasite *Toxoplasma* is the causative agent of toxoplasmosis, a worldwide foodborne zoonosis that is particularly severe when opportunistic or congenital. *Toxoplasma* has a complex life cycle that includes several distinct developmental stages (Extended Data Fig. 1a). The biology of the fast-growing tachyzoites and semi-dormant bradyzoites responsible for acute and chronic disease, respectively, is well known because they are easy to culture and benefit from well-established mouse models. In contrast, sexual reproduction is still largely *terra incognita*. Early studies in the 1970s using infected kittens elegantly but only partially documented the sexual cycle of *Toxoplasma* by scrutinizing the ultrastructure of the pre-gamete zoites and the sexual dimorphic stages in the intestinal lining of infected *Felis catus*^2–8^. All developmental stages have their own transcriptional signature, and switching between stages is controlled by intricate transcriptional cascades in which covalent and noncovalent epigenetic mechanisms act as driving forces^9, 10^. For example, tachyzoite and merozoite stages can be distinguished by their respective subtranscriptomes, with one being repressed during the other stage^11, 12^, a silencing function recently attributed to the chromatin modifier MORC. MORC functions as a central and early checkpoint of sexual commitment, and its conditional depletion promotes broad activation of sexual gene transcription in tachyzoites^1^. MORC forms a core complex with the histone deacetylase HDAC3, which bind strongly to a substantial number of Apetala (AP2) proteins^1^ (Extended Data Fig. 1b). AP2 have emerged as candidate transcription factors (TFs) in all apicomplexan species, in which they play a critical role in regulating life cycle transitions^9, 13^. It is thought that MORC/HDAC3-associated AP2s bind to specific sequences of DNA and direct the synchronized expression of stage-specific genetic programs^10^, but their relative contribution to *Toxoplasma* sexual fate still await further characterization. This study unveils the role of a complex network of transcriptional repressors in regulating the commitment to merogony in *Toxoplasma*.

**Fig. 1.**
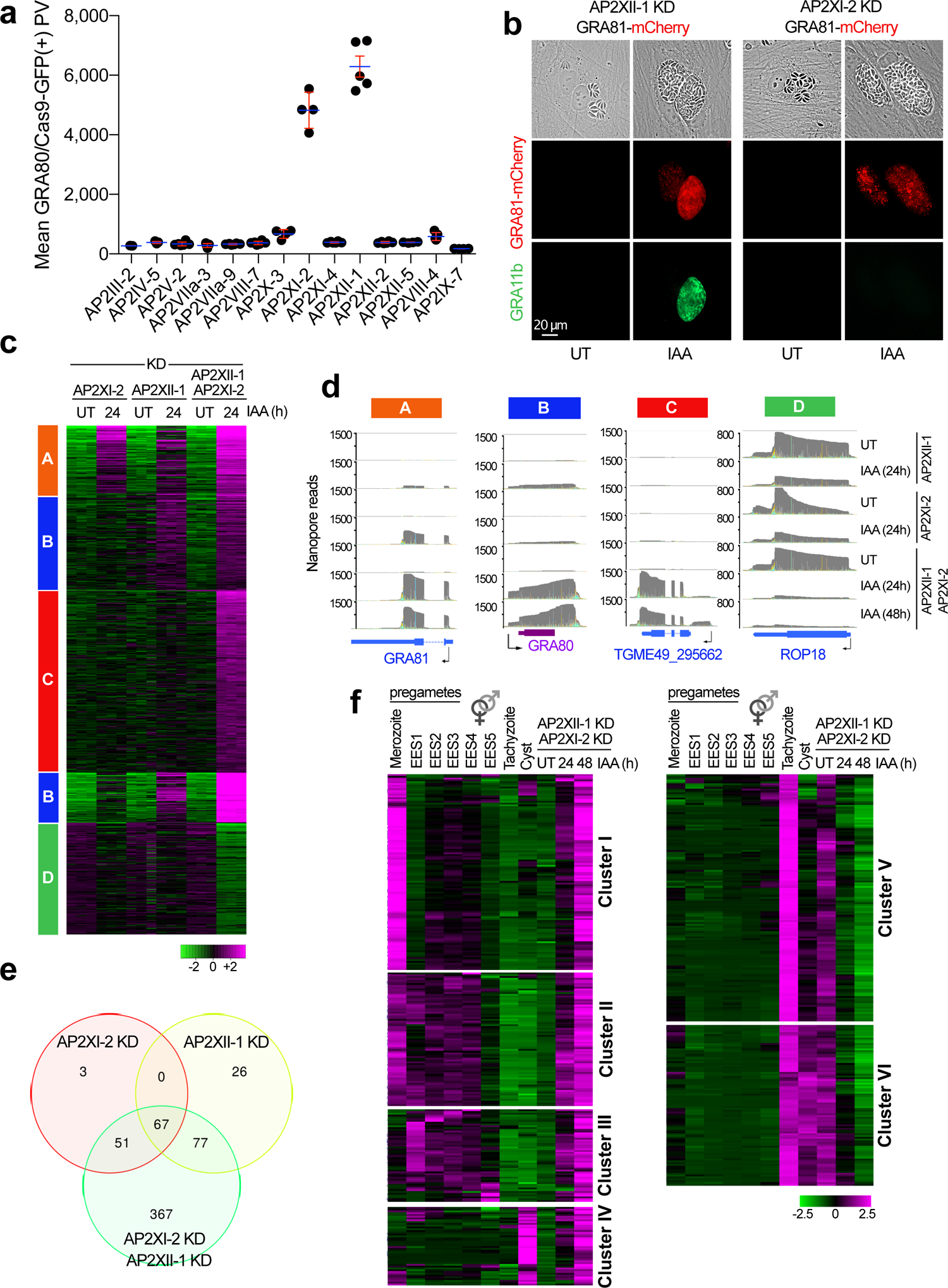
Simultaneous depletion of AP2XI-2 and AP2XII-1 induces the expression of merozoite-restricted transcripts. **a**, Expression of the merozoite marker GRA80 (*TGME49_273980*) was quantified *in situ* in intracellular zoites in which one of 14 MORC-associated AP2 was genetically disrupted. Cas9-GFP expression was used to assess the efficacy of genetic disruption in Cas9-expressing parasites (see Extended Data Fig. 1c). Horizontal bars represent the mean ± s.d. of GRA80 vacuolar intensity from three to four independent experiments (n = 50 GFP-positive vacuoles per dot). **b**, IFA of HFFs infected with parasites harboring a reporter gene (*TGME49_243940*) expressing GRA81, a merozoite protein endogenously tagged with mCherry within the RH AP2XII-1-mAID-HA or AP2XI-2-mAID-HA lineages. Untreated (UT) and IAA-treated zoites were probed with antibodies against GRA11b (green) and mCherry (red). **c**, Heat map of K-Means clustering (Pearson correlation) of 1500 variably expressed genes in three KD context. RPKM values were log2-transformed and mean centered then clustered using iDEP.96. Genes were grouped into four clusters on the basis of the expression similarity. **d**, M-pileup representation of aligned Nanopore reads at genes identified as up- or down-regulated in clusters A, B, C, or D after IAA-dependent knockdown of AP2XII-1 and AP2XI-2 individually or together. **e**, Venn diagram showing the overlap of genes that were upregulated in the three knockdown strains treated with IAA. **f**, Heat map of K-Means clustering (Pearson correlation) of 352 and 432 up- and down-regulated genes after simultaneous depletion of AP2XII-1 and AP2XI-2. RPKM values were log2-transformed and mean centered then clustered using iDEP.96 (Ge et al., 2018). Shown is the abundance of each transcript before and after depletion and during different life cycle stages, with data from transcriptomes of merozoites, longitudinal studies of enteroepithelial stages (EES1 to EES5), tachyzoites, and cysts published in ToxoDB.org. The color scale indicates log2-transformed fold changes.

## AP2XI-2 and AP2XII-1 jointly control gene expression

Using a loss-of-function CRISPR screen in *Toxoplasma*, we showed that of 14 MORC-associated AP2s, only AP2XI-2 or APXII-1, when inactivated, induced expression of GRA80 (*TGME49_273980*), a merozoite-specific protein to which we raised a specific antibody (Fig. 1a and Extended Data Fig. 1c). Because both AP2s are essential for tachyzoite growth, as indicated by their fitness scores (Extended Data Fig. 1b), we used the minimal auxin-inducible degron (mAID) system to acutely and reversibly deplete their protein levels and examine subsequent phenotypes. Proteins tagged with mAID undergo proteolytic degradation in the presence of indole-3-acetic acid (IAA) (Supplementary Fig. 1a, b). As expected, IAA treatment of these edited parasites resulted first, in a specific and complete degradation of the bulk of AP2XI-2-mAID-HA and APXII-1-mAID-HA proteins (Extended Data Fig. 1d) and second in a simultaneous accumulation of merozoite proteins at different expression levels, depending on the parasite line. Thus, the merozoite reporter gene GRA81-mCherry was induced upon depletion of each AP2 individually, whereas GRA11b, a hallmark of merozoites^14^, was expressed exclusively after loss of AP2XII-1, and GRA80 levels were less pronounced in AP2XI-2-depleted parasites (Fig. 1b and Extended Data Fig. 1e). Thus, the loss of AP2XI-2 does not fully mimic the depletion of AP2XII-1 in eliciting expression of the merozoite protein markers used here.

To examine the genome-wide transcriptional outcome of depleting each AP2 separately, we monitored mRNA levels using both Nanopore DRS and Illumina sequencing. After addition of IAA, substantial fractions of mRNAs were up- and down-regulated in a strain-specific manner, as shown by hierarchical clustering analyses (Fig. 1c and Supplementary Fig. 1c,d and Supplementary Table 2). AP2XI-2- and AP2XII-1-regulated subtranscriptomes could be defined by specific clusters (A and B, respectively, Fig. 1c) of genes induced as a consequence of their depletion, including *GRA80* and *GRA81* (Fig. 1d and Extended Data Fig. 1f). AP2XI-2 and AP2XII-1 also shared a small subset of differentially-regulated genes (n=67, Fig. 1e), suggesting that they function cooperatively or in the same pathways. We therefore examined the genetic relationship between these two TFs by editing a parasite strain with simultaneous knockdown (KD) of AP2XI-2 and AP2XII-1 compared with individual KD of either gene (Supplementary Fig. 1b). As expected, addition of IAA was accompanied by a near-complete loss of both proteins compared with untreated cells (Extended Data Fig. 1d). Double knockdown stimulated expression of genes from clusters A and B, but at a much higher level than single KD, and revealed a new cluster C (Fig. 1c,d and Extended Data Fig. 1f). Their depletion also resulted in suppression of gene expression (cluster D), a phenotype observed to a lesser extent with individual KD gene (Fig. 1c,d and Extended Data Fig. 1f). Simultaneous loss of both proteins resulted in a tailored transcriptional response characterized by more pronounced gene induction in terms of the number of genes affected and the expression levels of mRNAs and marked repression of a subset of genes, suggesting that AP2XI-2 and AP2XII-1 act synergistically or sequentially in regulating gene expression in *Toxoplasma*.

## AP2XI-2 and AP2XII-1 synergistically silence merozoite-primed sexual commitment

Comparative RNA-Seq analyzes showed that gene expression profiles after depletion of AP2XI-2 and AP2XII-1 perfectly matched the transcriptional state observed *in vivo* in enteroepithelial stages (EES) and merozoites collected from infected kittens^11, 12, 15^. Indeed, co-depletion of AP2XI-2 and AP2XII-1 predominantly and consistently induced the expression of genes specific to the pre-gametes at different developmental stages (clusters I, II, and III; Fig. 1f). A limited number of bradyzoite genes are also affected, but together they do not form a characteristic transcriptional signature associated with the latent stage (cluster IV; Fig. 1f). However, a subset of genes expressed exclusively in tachyzoites was simultaneously silenced (clusters V and VI, Fig. 1f), a trend also observed in merozoites from the cat intestine^11, 12^. The drastic changes in mRNA levels were also reflected in the abundance of the corresponding proteins: 18% of the 3,020 parasite proteins detected by mass spectrometry (MS)-based quantitative proteomics showed not only robust changes in abundance but also a highly polarized response to the merozoite stage, as underscored by our transcriptome analysis (Supplementary Fig. 2 and Table 3). In this context, the proteome of IAA-treated parasites shifted toward the pre-gametes stages, resulting in a significant over-expression of 276 proteins, whereas the expression of 285 tachyzoite proteins was suppressed after treatment (Supplementary Fig. 2c and Table 3). Treated parasites displayed RNA and protein expression profiles with a specific distribution of gene products (or functional categories), similar to enteroepithelial merozoites^11, 12^. Overall, housekeeping functions were not altered in IAA-treated parasites as confirmed by our proteomic analysis, in which no significant changes were detected in 82% of the 3,020 quantified proteins. Nor was their metabolic capacity changed, with the exception of induction of genes involved in purine metabolism, a phenotypic feature of merozoites found in the gut of cats^12^.

## Simultaneous depletion of AP2XI-2 and AP2XII-1 switches from tachyzoites to merozoites features

The process of invasion has been thoroughly examined, revealing the cryptic functions of organelle-resident proteins, specifically in the tachyzoite stage^16^. First, micronemes secrete proteins (MICs) that function as adhesins that facilitate attachment to the host cell and thus play a key role in motility and invasion. Following attachment to host cells, rhoptries release their neck (RON) and bulb (ROP) proteins, which interact with MIC proteins and contribute to parasite entry by opening the host cell membrane and directing the formation of parasitophorous vacuole. This function is primarily attributed to the RON complex. The final wave of secretion occurs with the release of proteins from the dense granules (GRA) that are involved in intravacuolar function, such as the formation of a tubulovesicular network (TVN), but also act at the PVM or operate as extravacuolar effectors to subvert host signaling pathways and reprogram host gene expression^17^. It was reported that the majority of MIC, ROP, and GRA proteins secreted by tachyzoites and bradyzoites are not expressed in merozoites^11, 12, 18, 19^. Consistently, we observed a complete switch in the expression of many known MIC, ROP, and GRA proteins upon addition of IAA as well as undescribed secretory proteins that appear to be involved in promoting asexual replication of merozoites (Supplementary Fig. 3).

For example, in IAA-treated parasites, MIC proteins that are highly expressed in tachyzoites and secreted as functional complexes, were strongly suppressed (e.g., MIC2 and AMA1; Fig. 2a). Conversely, merozoite-specific MICs such as the MIC17a,b,c cluster and AMA2, the ortholog of AMA1 in pre-gametes, were markedly induced (Fig. 2a), and transcriptional reprogramming was highly specific, as levels of SporoAMA1, its counterpart in the sporozoite, a quiescent stage found in sporulated oocysts, remained unchanged (Supplementary Table 2).

**Fig. 2.**
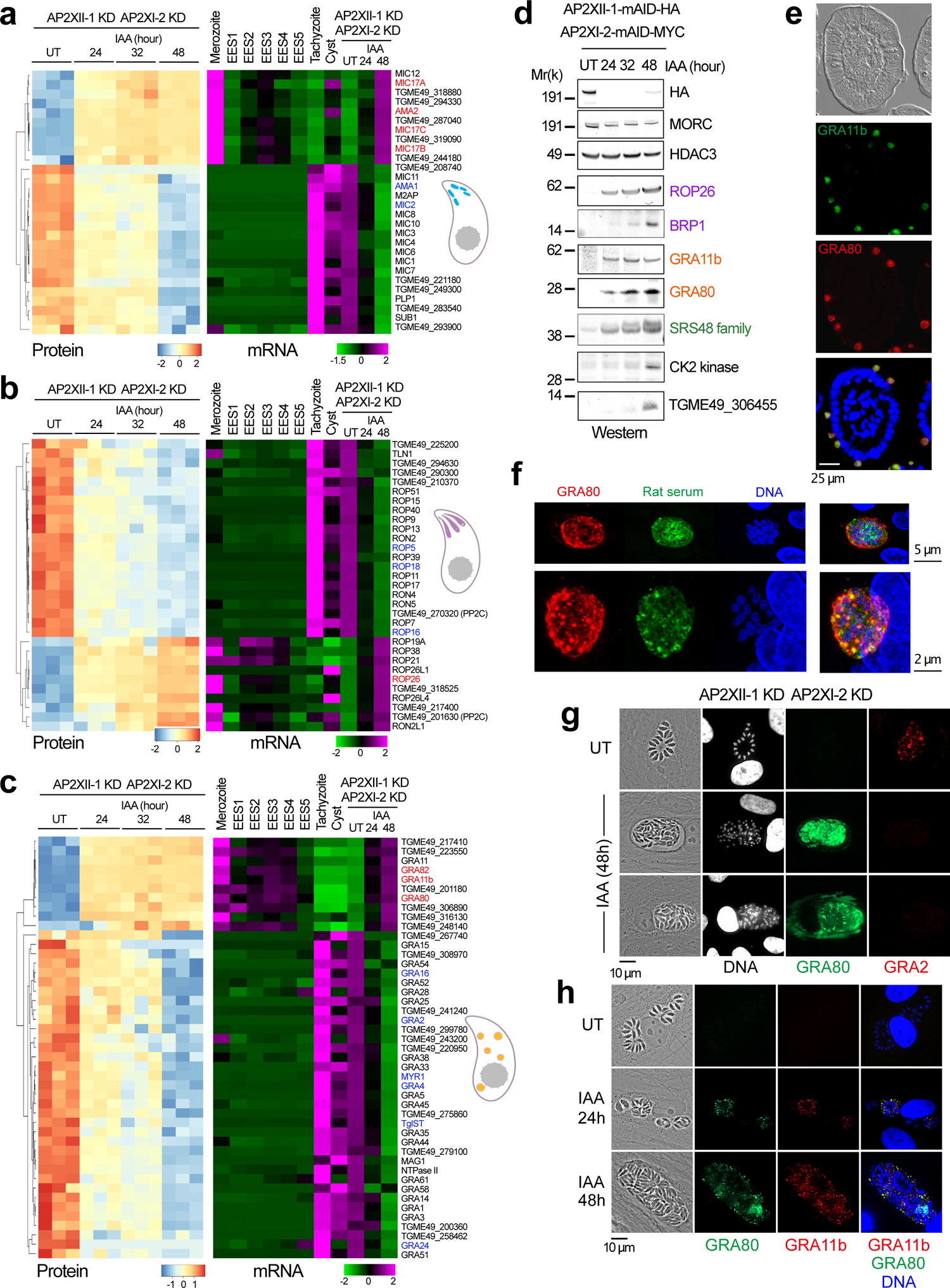
Co-depletion of AP2XI-2 and AP2XII-1 causes rewiring of organellar proteomes specialized in invasion and host-parasite interaction. **a-c**, Heat map showing hierarchical clustering analysis (Pearson correlation) of selected rhoptry (**a**), dense granule (**b**) and microneme (**c**) mRNA transcripts and their corresponding proteins differentially regulated after simultaneous and conditional depletion of AP2XII-1 and AP2XI-2. Shown is the abundance of their transcripts at different developmental stages, namely tachyzoite, cyst, merozoite, and EES. The color scale indicates the log2-transformed fold changes. The genes of interest are highlighted in red. **d**, Time-course western blot analysis of protein expression levels after depletion of AP2XII-1mAID-HA and AP2XI-2-mAID-MYC. Samples were collected at the indicated time points after addition of IAA and probed with antibodies against HA, MORC, and HDAC3, rhoptry proteins (ROP26 and BRP1), dense granule proteins (GRA11b and GRA80), the SRS48 family, a CK2 kinase (*TGME49_307640*), and the protein encoded by *TGME49_306455*. The experiment was repeated three times and a representative blot is shown. **e**, Epifluorescence image of IFA of infected small intestine of kittens stained with antibodies against GRA80 (red) and rat immune serum (green). Nuclei were counterstained with DAPI. **f**, Maximal intensity projection of confocal stacks after IFA with antibodies against GRA80 (red) and rat immune serum (green). Nuclei were counterstained with DAPI. **g**, Expression of tachyzoite protein GRA2 (red) and merozoite protein GRA80 (green) after knockdown of AP2XII-1 and AP2XI-2 was measured by IFA. **h**, Expression of merozoite proteins GRA11b (red) and GRA80 (green) was measured by IFA 24 and 48 hours after addition of IAA. Cells were co-stained with Hoechst DNA-specific dye.

The combined depletion of AP2XI-2 and AP2XII-1 silenced 80% of the 143 rhoptry proteins described or predicted by hyperLOPIT to be tachyzoite-specific (Fig. 2b and Supplementary Fig. 3a). These include the components of the RON complex as well as ROP16, ROP18, and ROP5, which function as effectors to protect parasites from host cellular defenses and thwart immune responses. Only a few known rhoptry proteins were induced, including BRP1, which is abundant in both bradyzoite and merozoite stages^20^, and a family of ROP5-related kinases, referred to here as the ROP26 family, which is expressed exclusively in pre-gametes (Fig. 2d).

We observed a similar tendency toward silencing of the tachyzoite program and activation of the merozoite program when we examined GRA mRNA and protein levels in response to co-depletion of AP2XI-2 and AP2XII-1 (Fig. 2c and Supplementary Fig. 3b). For example, the levels of the TVN core proteins, i.e. GRA2, GRA4, and GRA6, decreased dramatically over time, as did the MYR-dependent effectors GRA16, GRA24, or TgIST in IAA-treated parasites (Fig. 2c). In striking contrast, genuine pre-gametes markers (GRA11a and GRA11b^14^), as well as unannotated set of GRA proteins, were expressed exclusively in merozoites (Fig. 2c), such as GRA80, GRA81, and GRA82 showing a punctate pattern in the IAA-treated parasites that occasionally overlap with the canonical merozoite GRA11b (Fig. 2g-h; Extended Data Fig. 1e and Fig. 2a). Upon secretion, GRA80, GRA81, and GRA82 localized to the vacuolar space and/or the PV membrane (PVM), but differ in their secretion kinetics. Interestingly, GRA80 labelled mature schizonts that reproduce asexually in the cat small intestine (Fig. 2e, f). GRA80 was detected *in vitro* in dense granules at an early time point after addition of IAA, then secreted in the vacuolar space to associate with the PV membrane both in cat gut stages (Fig. 2f) and *in vitro* in cultured fibroblasts (Fig. 2g,h). GRA80 eventually crossed the PVM to spread into the host cell cytoplasm (lower panel in Fig. 2g). On the other hand, GRA82 was detectable only after 48 hours in the mature merozoite and contrasts with the kinetics of GRA11b and GRA80, which are induced in the parasite and the vacuole at the onset of merogony (Extended Data Fig. 2a).

This transcriptome/proteome shift from the tachyzoite to the merozoite program also occurs in other protein families and leads, for example, to a dramatic restructuring of the proteins on the surface of the zoite after IAA treatment, including the deemed SAG-Related Surface (SRS) protein family (Extended Data Fig. 2b). Compared to tachyzoites, merozoites express the largest repertoire of SRS, e.g., the SRS48 (Fig. 2d) and SRS59 (Extended Data Fig. 2c) families, which have been predicted to promote gamete development and fertilization^12^. Accordingly, 90% of the known 88 SRS were induced in IAA-treated parasites phenocopying the merozoite stage, while all tachyzoite-specific SRS were simultaneously suppressed (Supplementary Fig. 3d). This transition to pre-gametes is also accompanied by the expression of 29 of the 33 members of Family A in treated parasites (Extended Data Fig. 2d,e), which are recognized as predominant secreted and/or membrane-associated merozoite proteins^11, 12^.

Overall, AP2XI-2- and AP2XII-1-depleted parasites share common features with *bona fide* merozoites, notably the abrogation of the characteristic molecular signature of tachyzoites and expression of a specialized merozoite proteome. Phenotypically, treatment with IAA results in a decrease in parasite infectivity over time, likely due to suppression of key proteins required for motility, attachment, or invasion in tachyzoites. As a result, *in vitro* converted merozoites fail to egress and to re-invade new host cells after a period of proliferation, as evidenced by the dramatic reduction of lytic plaques in treated parasites compared to untreated parasites (Extended Data Fig. 2f).

## AP2XI-2/AP2XII-1-depleted parasites undergo several rounds of schizogonic replication to produce merozoites

Our understanding of pre-sexual stages at the cellular level goes back to the original description by Dubey and Frenkel (1972)^21^, who systematically recorded the morphological details of five pre-gamete stages, designated merozoites A to E, proposed to be formed sequentially during colonization of the epithelial cells of the cat intestine prior to gamete formation (Fig. 3a). These morphotypes vary in size and shape, appearing in different areas of the cat’s digestive tract asynchronously, making challenging the analysis of pre-sexual stages *in vivo*^18, 19, 22–24^. We then investigated to which extent the zoites produced during *in vitro* induced merogony share the same morphological structural features as their counterparts observed *in vivo*. For this purpose, we performed IFA using antibody toolkit originally developed for studying the subcellular content and division of tachyzoites (i.e., antibodies recognizing inner membrane complex or IMC^25^) and transmission electron microscopy (TEM).

**Fig. 3.**
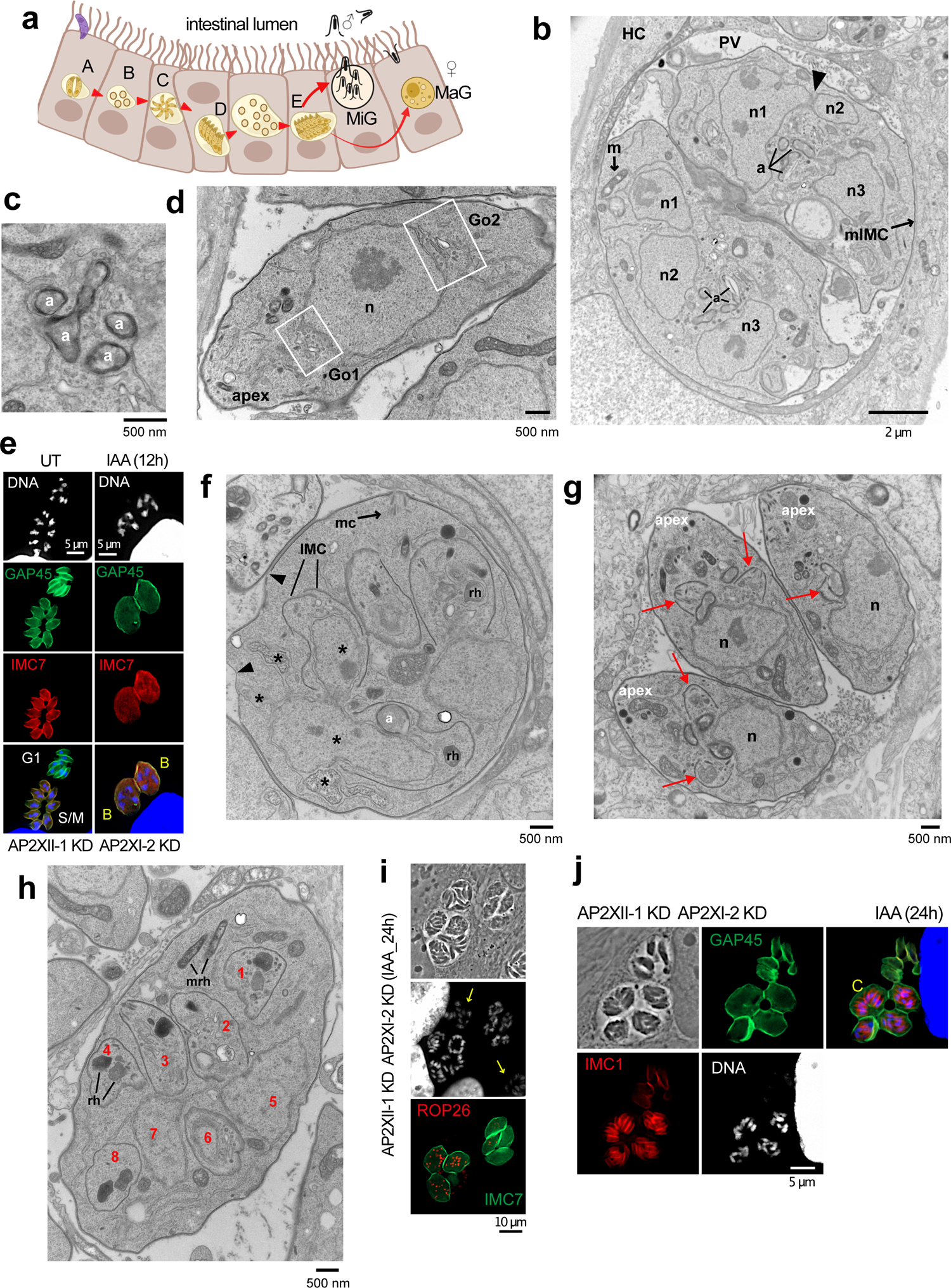
AP2XI-2/AP2XII-1-depleted zoites undergo endopolygeny with karyokinesis. **a**, Cartoon showing the merogony process along the intestinal tract. Bradyzoites (in pink) sequentially differentiate into merozoites A-E before giving rise to macro-gametes (MaG) and micro-gametes (MiG). Electron micrograph images of RH (AP2XII-1 KD/AP2XI-2 KD)-infected HFFs untreated (**g**) or treated for 24 hours with IAA (**b-d**, **f**, **h**). **b**, Emphasis on karyokinesis with fission. n1 to n3: nuclear profiles, a: apicoplast, m: mitochondrion, mIMC: mother inner membrane complex, hcell: host cell, PV: parasitophorous vacuole. Arrowhead shows nuclear fission. **c**, Emphasis on apicoplast multiplication by growth and scission. **d**, Emphasis on Golgi multiplication from either side of the nucleus. Go1 and Go2: 2 Golgi apparatus. **e**, IFA of tachyzoites (UT) and AP2XII-1/AP2XI-2-depleted zoites (12 hours post-IAA). GAP45 stains the mother cell and its progeny. IMC7 specifically stains the diploid (left) and polyploid (right) mother cell. Cells were stained with Hoechst DNA-specific dye. Types B, meronts are marked in yellow. **f**, Emphasis on appearance and role of the IMC segregating daughter buds in the mother cytoplasm. mc: mother conoid, a: apicoplast, IMC: inner membrane complex, rh: rhoptry. Arrowhead shows area devoid of the IMC and asterisks highlight the nuclear fissions. **g**, Emphasis on contrasting endodyogeny in tachyzoites (untreated condition). Two daughter buds formed apically and symmetrically (arrows). **h**, Emphasis on endopolygeny showing up to 8 daughters buds and ultrastructure of rhoptries. rh: rhoptry, mrh: mother rhoptry. **i-j**, AP2XII-1/AP2XI-2-depleted meronts (24 hours post-IAA) were fixed and stained with ROP26 (*TGME49_209985*) or IMC1 (red), IMC7 (green) and Hoechst DNA-specific dye (white or blue). Yellow arrows indicate IMC7-negative mature merozoites and type C meronts are shown.

As a first step 24h post-IAA addition, the nucleus of the mother cell undergoes multiple fissions with the maintenance of the nuclear envelope, leading to individualized nuclei (in even numbers, varying from 4 up to 10) (Fig. 3b). Concomitantly, single organelles, such as the apicoplast and the Golgi apparatus expand and multiply to reach number equal to the number of the nucleus. Transversal cross-sections of the apicoplast (limited by four membranes) reveals its elongation and constriction, suggestive of replication by scission (Fig. 3c), which was also visualized by immunofluorescence with the ATrx1 antibody^26^ (Extended Data Fig. 3a) and is consistent with maternal inheritance of the apicoplast observed in meronts in the cat intestine^27^. Multiple Golgi complexes are formed at different sites of the nucleus, sometimes in opposing orientations, suggestive of *de novo* formation (Fig. 3d); the multiplication mode of other organelles, like secretory organelles are unknown. At this stage, the multinucleated mother cell contains several sets of organelles randomly distributed throughout the cytoplasm. Despite the increase in size of the mother cell, the subpellicular IMC is still prominently present beneath the plasma membrane. The parasites that exhibit a characteristic ovoid shape with 4 and 8 nuclei are morphologically related to the cryptic and early meronts, namely B and C morphotypes^21^ (Fig. 3e and Extended Data Fig. 3b).

As a second step, new flattened vesicles of the IMC appear in the mother cytoplasm, and progressively elongate allowing the sub-compartmentalization of organelles destinated for each daughter cell (Fig. 3f). This process of internal budding of more than two daughter cells (referred here as endopolygeny)^23, 28–31^ differs from the tachyzoite division by endodyogeny in which the two daughter cells are generated symmetrically and in a synchronous manner in the mother cell (Fig. 3g). Alongside with the expansion of daughter buds, the mother IMC and conoid show partial disassembly. Interestingly, rhoptries inside daughter cells are different in shape and electron-density from mother rhoptries dispersed in the cytoplasm, suggesting *de novo* biogenesis of rhoptries (Fig. 3h) which can be also followed with the merozoite-specific protein ROP26 (Fig. 3i). This finding is in line with the observation that the bulbous end of the rhoptry in the meronts of infected cats remains spherical, in contrast to tachyzoites and bradyzoites^24^. In these multinucleated structures, it was possible to identify daughter IMC stained with IMC1, whereas GAP45 staining was restricted to the periphery of the mother cell (Fig. 3j, Extended Data Fig. 3c). Daughter cells become polarized with the formation of a conoid and apical distribution of micronemes, rhoptries, the apicoplast and the Golgi apparatus (Fig. 4a, Extended Data Fig. 3d). At this stage, it is clear that the maternal conoid coexists with the newly formed conoids of the progeny, as shown by the labelling of apically methylated proteins (Fig. 4b). After final assembly, the daughter cells emerge separately, wrapped by the plasma membrane of the mother cell (cortical or peripheral budding) (Fig. 4c,d), forming fan-like structures as previously described in infected cat cells^23^.

**Fig. 4.**
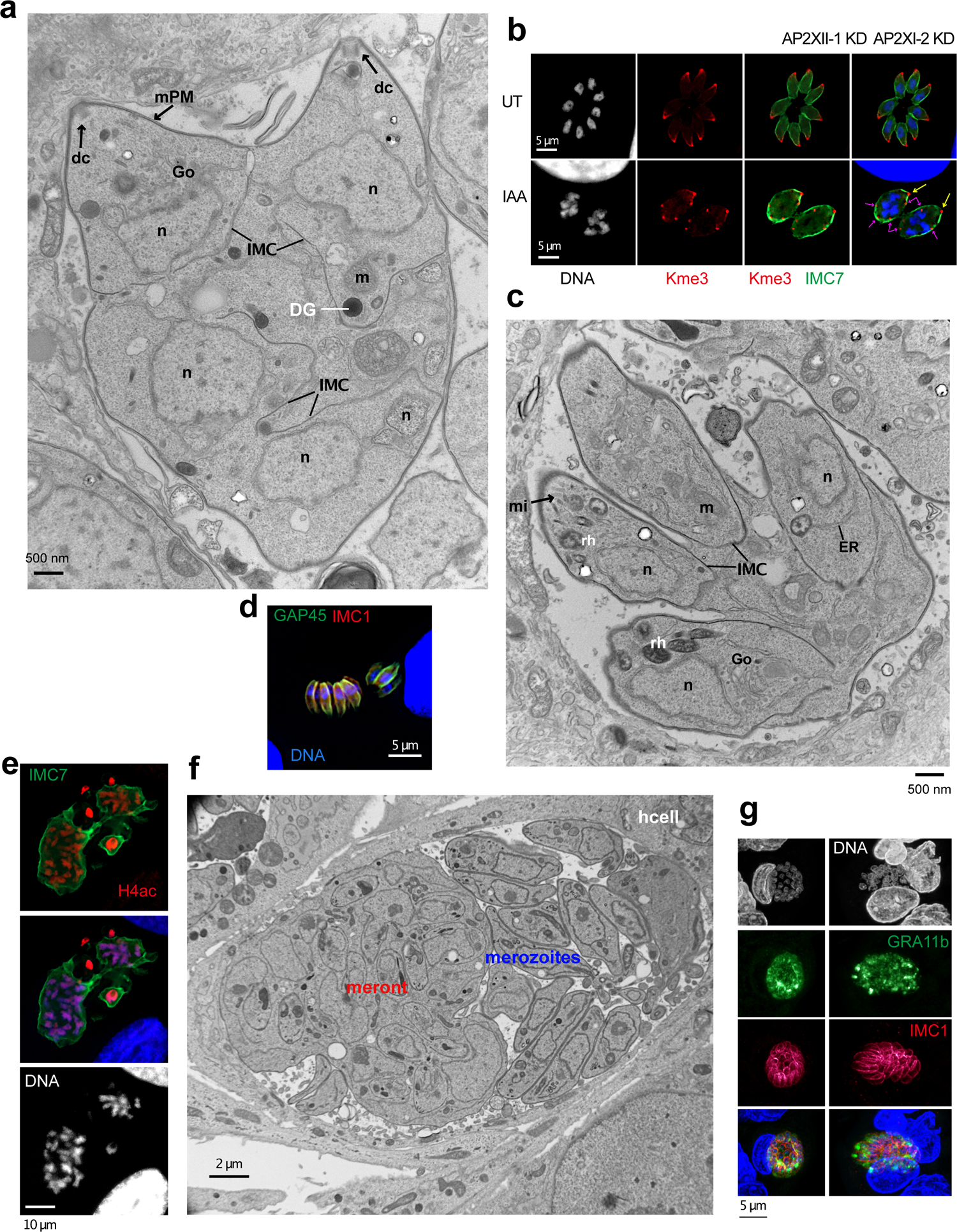
*In vitro* induced merogony typified by the emergence of multiple zoite stages. Electron micrograph images of RH (AP2XII-1 KD/AP2XI-2 KD)-infected HFFs treated for 24 hours (**a, c**) or 48 hours with IAA (**f**). **a**, Emphasis on protruding daughters sharing the mother plasma membrane. mPM: mother plasma membrane, Go: Golgi apparatus, n: nucleus, DG: dense granule, m: mitochondrion, dc: daughter conoid. **b**, Tachyzoites (UT) and AP2XII-1/AP2XI-2-depleted meronts (24 hours post-IAA) were fixed and stained with H3K9me3 (red) and IMC7 (green) and Hoechst DNA-specific dye (white or blue). Yellow and pink arrows indicate mother and daughter conoids, respectively. **c**, Emphasis on daughter cell emergence. mi: microneme, m: mitochondrion, ER: endoplasmic reticulum, Go: Golgi apparatus, n: nucleus, rh: rhoptry, IMC: inner membrane complex. **d,** Representative image of neatly aligned elongated merozoites and forming fan-like structures as they hatch from the mother cell. Mature merozoites are co-stained with GAP45 (green), IMC1 (red) Hoechst DNA-specific dye (blue). **e**, Image of a giant schizont delineated by IMC7 (green) showing polyploidy (n=16). The nuclear structure is co-stained with Hoechst DNA-specific dye and hyperacetylated histone H4 (red). **f**, Emphasis on large PV containing a mega meront with many daughter buds residing with merozoites in the same PV. hcell: host cell. **g**, Maximal intensity projection of a confocal microscopy z-stack from meront in infected small intestine of a kitten. Antibodies against GRA11b (green) mark the dense granules and against IMC1 (green) the inner membrane complex of individual merozoites. Nuclei are counterstained with DAPI.

Compared to tachyzoites, these newly formed parasites are thinner and do not form a rosette-like structure within the PV but instead are aligned, with their apex facing the PV membrane (Extended Data Fig. 3e,f); these features are reminiscent to those of type D-like merozoites produced in feline intestinal cells from meront entities at the onset of infection^29–31^. Interestingly, the PV membrane of merozoites also forms physical interactions with host ER and mitochondria, perhaps for nutrient acquisition^32^ (Extended Data Fig. 3e). A frequent observation is the presence of multiple parasites in a single PV undergoing endopolygeny (Supplementary Fig. 4a,b). When we monitored the degree of ploidy with the DNA-specific dye Hoechst and by staining with a centrosome marker (human centrin, Supplementary Fig. 4c,d) or pan-acetylated histone H4 (Supplementary Fig. 4e), we observed that nuclear division cycles within the same schizont were not synchronous.

## Fully developed merozoites produced *in vitro* have conserved and distinct subcellular features

Extension of the IAA treatment for additional 16 hours reveals the presence of very large meronts containing numerous daughter cells in formation (Fig. 4e,f). These meronts are detected in the same PV together with fully formed merozoites (Fig. 4f), as their counterpart in the cat gut (Fig. 4g). Mature polyploid meronts can be visualized by IMC7 staining on their surface, whereas fully formed merozoites were completely negative for IMC7 (Fig. 3e,i, Extended Data Fig. 3b), a phenotype also been observed in pre-gametes developing in the cat gut^25^. Remarkably, we identified new merozoite-specific markers that clearly distinguish the two morphotypic populations. For example, ROP26 exclusively marks zoites undergoing schizogonic replication, in contrast to GRA11b and GRA80 expression, which are restricted to mature merozoites (Fig. 3i and Extended Data Fig. 4a). As merozoites undergo several cycles of endopolygeny, they acquire novel distinct morphological features compared to first-generation merozoites (24 hours post-IAA), likely type E^23^. Some merozoites appear sausage-shaped, with a diameter of 1.5-1.8 micron, packed in the PV without any spatial organization (Extended Data Fig. 4b,c). These forms contain similar organelles found in tachyzoites but surprisingly, they exhibit an extruded conoid (Extended Data Fig. 4d). Other PV contain peripherally arranged parasites, leaving a large empty space (Extended Data Fig. 4e,f), reminiscent to schizont PV formed in feline intestinal cells^23^. Interestingly, these parasites at the PV edge adopt two configurations: either they have a very large cell body (trapezoid) with a diameter up to 5 microns or a very thin and elongated shape (tubular), with a diameter of 200-250 nm (Extended Data Fig. 4g,h). These latter do not contain nucleus but mitochondria profiles and ribosomes are observed. Their origin and formation remain to be determined but their abundance in PV likely suggest a physiological relevance in the *Toxoplasma* lifecycle.

## AP2XI-2 and AP2XII-1 bind as a heterodimer to HDAC3 and MORC

AP2XI-2 and AP2XII-1 likely synergize to suppress gene expression in tachyzoites, but their *modus operandi* is still enigmatic. Both proteins were originally found in a MORC pulldown along with HDAC3 in tachyzoites^1^. We confirmed their strong and specific association with MORC/HDAC3 by reverse immunoprecipitation combined with MS-based proteomic and Western blot analyses using knock-in parasite lines expressing a FLAG tagged version of AP2XI-2 or AP2XII-1 (Fig. 5a,b). MS-based proteomics found both AP2 proteins in co-immunoprecipitation eluates, indicating that they are part of the same operating complex (Fig. 5c and Supplementary Table 4 and 5). To support this hypothesis, we used baculovirus to transiently co-express epitope tagged AP2XI-2-Flag and AP2XII-1-(Strep)2 in insect cells (Fig. 5d). AP2XII-1 was purified by Strep-Tactin affinity chromatography and its partnership with AP2XI-2 was confirmed by Western blotting and MS-based proteomics (Fig. 5e and Supplementary Table 6). Consistent with AP2XI-2 and AP2XII-1 being part of a heterodimer, these two proteins coelute in the same gel filtration fractions, in a MORC- and HDAC3-independent manner (Fig. 5e). Many transcription factors, including apicomplexan AP2 were reported to form homo- and heterodimers with different partners that modulate DNA binding specificity and affinity^33, 34^. In this context, AP2XI-2 and AP2XII-1 likely bind cooperatively as a heterodimer to DNA to selectively and synergistically repress merozoite gene expression, and only their simultaneous depletion leads to achievement of the developmental program critical for merozoite formation.

**Fig. 5.**
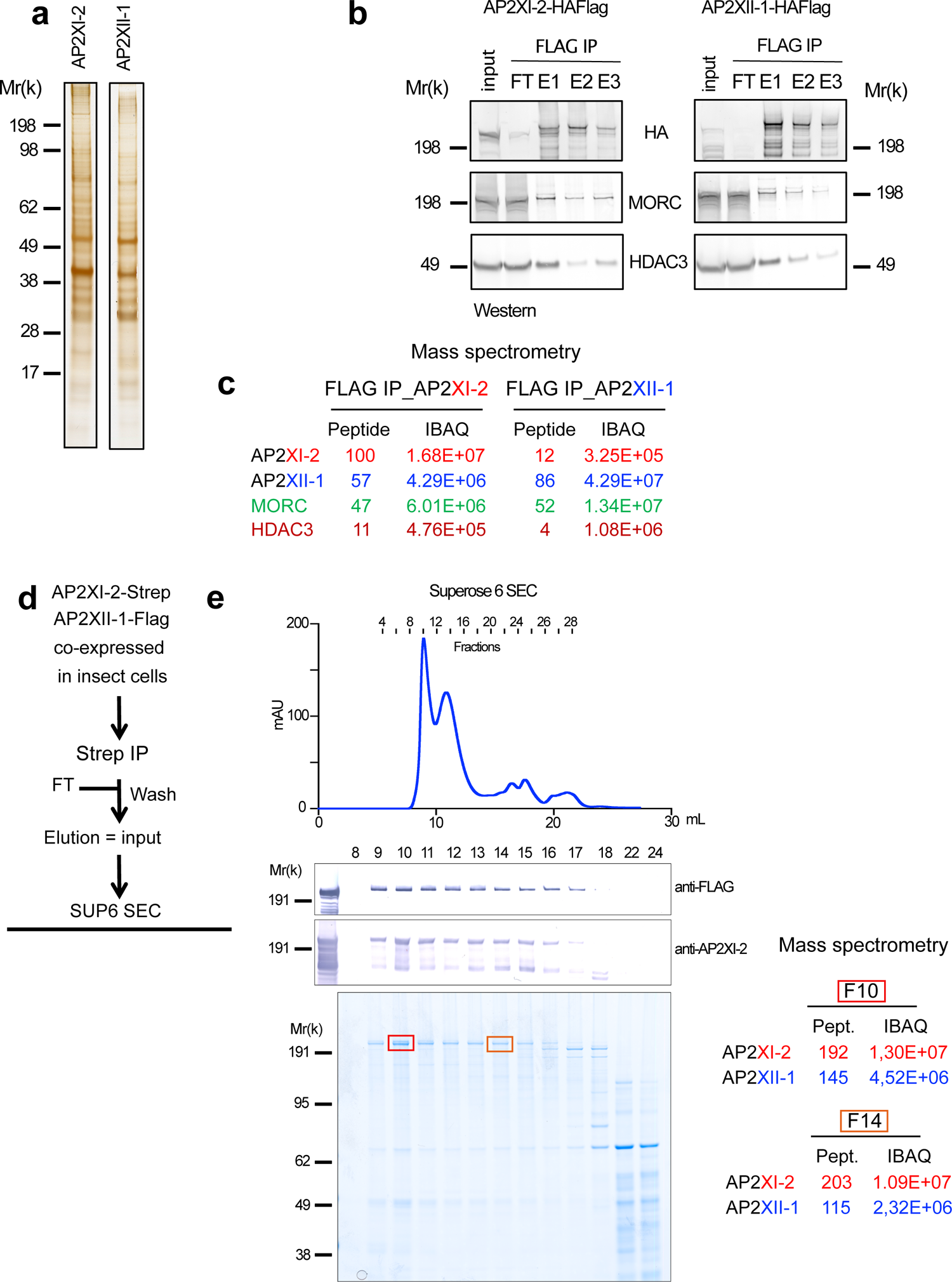
AP2XII-1 and AP2XI-2 heterodimerize and interact with HDAC3 and MORC to form a repressive core complex. **a**, AP2XII-1 or AP2XI-2 were ectopically tagged with HAFlag in the RH strain, and their associated proteins were purified by FLAG chromatography. Silver staining shows their first eluted fraction. **b**, Flag affinity eluates were analysed by Western blot to detect MORC and HDAC3. **c**, MS-based proteomic analysis of AP2XI-2 and APXII-1 FLAG elutions identified MORC and HDAC3 subunits and APXII-1 in the AP2XI-2 purification and *vice versa*. Number of identified peptides and intensity based absolute quantification (iBAQ) value are indicated. **d**, Chromatographic purification scheme of AP2XI-2-Strep and APXII-1-Flag co-expressed in *Trichoplusia ni* (Hi-5) insect cells. **e**, Strep-tactin XT purified proteins were fractionated on a Superose-6 increase gel filtration column. Input (strep-tactin elution) and gel filtration fractions were separated by SDS -polyacrylamide gel and analyzed by Western blot using anti FLAG and in-house anti-AP2XII-2 antibodies. Fraction numbers are indicated at the top of the gel. MS-based proteomic analyzes of fractions 10 and 14 are indicated on the right side of the graph. Number of identified peptides and iBAQ values are indicated.

## AP2XI-2 and AP2XII-1 colocalize extensively in the genome where they recruit MORC and HDAC3 to reduce chromatin accessibility

To further explore how AP2XI-2 or AP2XII-1 cooperate in *Toxoplasma* to silence gene expression, we examined their genome-wide distribution using chromatin immunoprecipitation followed by deep sequencing (ChIP-seq; GSE222819) in the context of their conditional single or double knockdown. In parallel, we generated high-resolution profiles of MORC and HDAC3 using in-house ChIP-grade antibodies. The enrichment at chromatin of AP2XII-1, which is the highest, was used as a proxy to analyze the distribution of loci to which the repressive complex binds. As a result, we identified two clusters of genes with significant enrichment in their environment: Cluster 1 groups genes with a scattered distribution of peaks that span large chromosomal regions and have low enrichment, making them less likely targets. In contrast, genes in Cluster 2 exhibit a discrete peak with a high amplitude centered at known or predicted transcription start site (TSS) (Extended data: Fig. 5a). Interestingly, integrative analysis of RNA-seq and ChIP-seq data showed that genes from Cluster 2 are exclusively expressed in the pre-gametes and as such match well with genes from Cluster I to IV mined from transcriptome data (Fig. 1f).

To investigate the co-occupancy of AP2XII-1 and its partners, we focused our analysis on Cluster 2 genes. As predicted, when IAA was added, there was a nearly complete loss of AP2XI-2 and AP2XII-1 chromatin enrichment compared to the untreated parasites (Fig. 6a). Overall, the inspection of these individual ChIP–seq tracks revealed a strong overlap between the binding sites of the AP2XI-2 or AP2XII-1 cistromes (Fig. 6b and Extended Data Fig. 5b), with approximately 30-50% of the peaks located at the TSS (Extended Data Fig. 5c). AP2XII-I and AP2XI-2 showed similar genome-wide occupancy when immunoprecipitated from single or double knockouts (Extended Data Fig. 5d-e, h). We next investigated whether AP2XI-2 and AP2XII-1 are individually or jointly required for the recruitment of MORC and HDAC3 to chromatin by examining their genome-wide occupancy in the absence of AP2XI-2 and/or AP2XII-1. After the addition of IAA and acute depletion of these AP2s, we observed a concomitant reduction in HDAC3 and MORC occupancy at TSS of cluster 2 genes, which is even more pronounced in the context of double knockdown (Fig. 6a; Extended Data Fig. 5h).

**Fig. 6.**
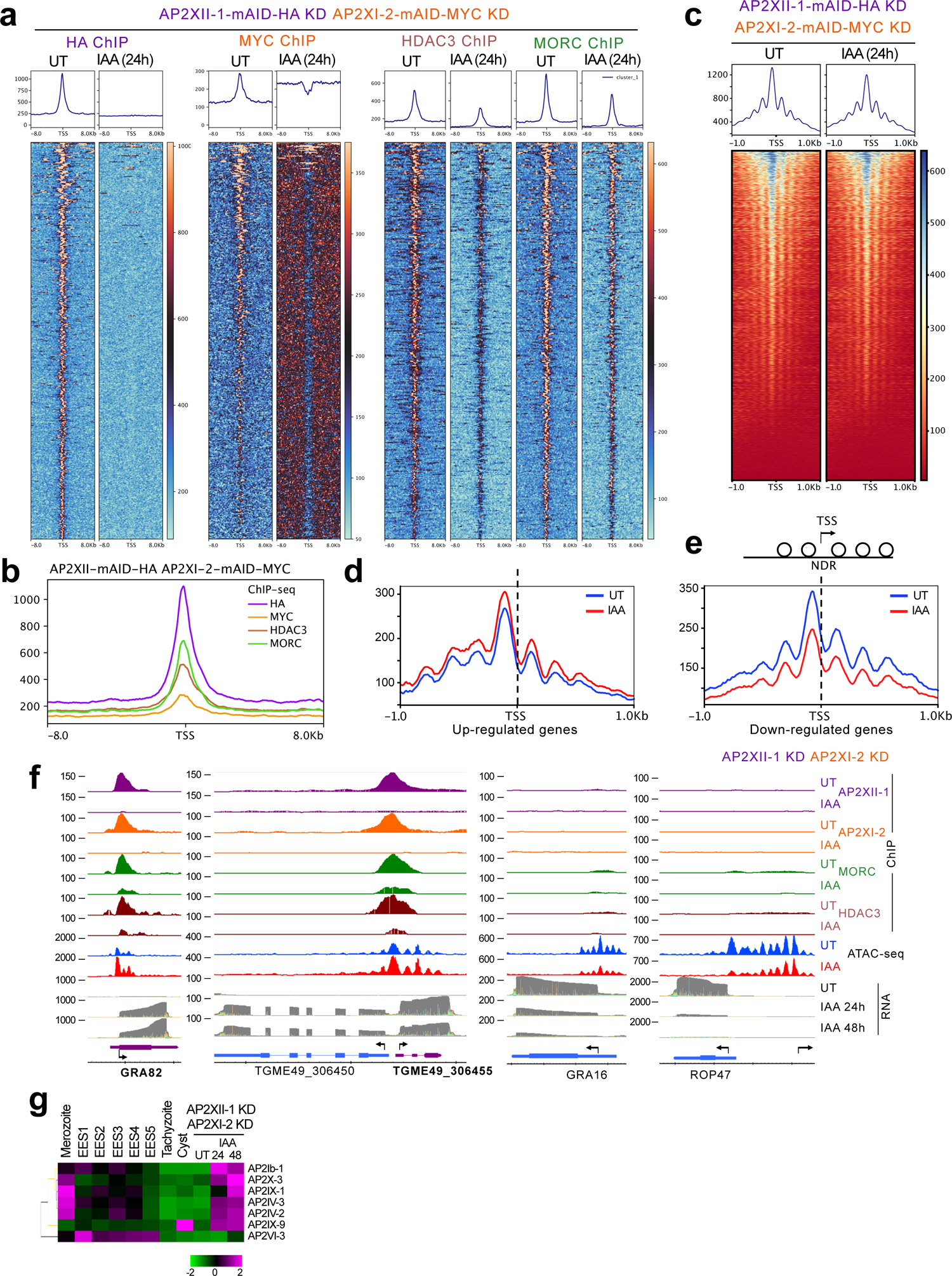
Recruitment of HDAC3 and MORC to chromatin is mediated jointly by AP2XII-1 and AP2XI-2. **a**, Profile and heat maps of averaged sum ChIP-seq called peaks showing binding of APXII-1 (HA), AP2XI-2 (MYC), HDAC3, and MORC in the vicinity of TSS of annotated gene promoters before and after addition of IAA. The top panels show the average signal profile on genomic loci centered at TSS (±8 kb). The lower panels show heat maps of peak density around the same genomic loci. The color scale for interpreting signal intensity is on the right side of each graph. **b**, Superposed profile plots show the co-enrichment of APXII-1 (HA) ChIP-Seq signals with AP2XI-2 (MYC), HDAC3, and MORC signals around TSS of annotated gene promoters in the untreated state. **c**, Profile and heat maps of averaged sum show Tn5 transposase accessibility (ATAC-sequencing) for *T. gondii* genes centered at TSS (±1 kb) in untreated (UT) and IAA-treated (24 hours) samples. High read intensity is shown in blue. The average signal profile is plotted above. (**d**, **e**) Tn5 Transposase accessibility plot in *T. gondii* allows for nucleosomal occupancy prediction. Comparison of coverage profiles of ATAC-seq read signals of untreated and IAA-treated (24 hours) across down-regulated genes (n=226) and up-regulated genes (n=281) relative to TSS (±1 kb). ATAC-seq enrichment plot shows that the nucleosome-depleted region (NDR) at TSS, while mono-nucleosome fragments are enriched at flanking regions and show phased nucleosomes at the −2, −1, +1, +2, and +3 positions. **f**, IGB screenshots of four genomic regions with representative merozoite and tachyzoite genes. ChIP-seq signal occupancy for HA, MYC, MORC, and HDAC3 are shown in the untreated state and after simultaneous depletion of AP2XII-1-mAID-HA and AP2XI-2-mAID-MYC. RNA-seq data for different induction time points are shown as M-pileup representation of the aligned nanopore reads. Tn5 Transposase accessibility profiles are plotted for both conditions. **g**, Heat map showing hierarchical mRNA clustering analysis (Pearson correlation) of AP2 TFs regulated by simultaneous depletion of AP2XII-1 and AP2XI-2. Shown is the abundance of their transcripts at different developmental stages, namely merozoites, EES, tachyzoites, and cysts. The color scale indicates the log2-transformed fold changes. Heatmap and profile plots (panels **a**-**e**) were generated using the Deeptools program. Panels **a, b** use the Extended Data Fig 5a cluster 2 geneset while panel **d** and **e** use up- and down-regulated genesets established by Illumina RNA sequencing.

AP2XI-2 and AP2XII-1 are expected to mediate chromatin compaction and accessibility, a function attributed to their partners MORC and HDAC3. To investigate this assumption, we performed ATAC-seq (GSE222832), a robust and streamlined method for profiling chromatin accessibility^35^. ATAC-seq peaks represent regions of chromatin accessible to transposases and are therefore proxies for transcription factor occupancy on chromatin. At the genome level, there is a slight decrease in average accessibility between untreated and treated conditions (Fig. 6c) with a majority of the peaks located at TSS (Extended Data Fig. 6a,b). However, when we plotted ATAC-seq data for the subset of down- and up-regulated genes (as defined in Fig. 1f), the changes in occupancy were more pronounced and coherent with expected increase or decrease in accessibility of induced or repressed clusters, respectively (Fig. 6d,e). Additionally, in many *Toxoplasma* promoters, we observed that two well-positioned nucleosomes upstream and downstream of TSS define a nucleosome-depleted region (NDR) that serves as a binding platform for RNA polymerase and transcription factors (Fig. 6e).

At the gene level, and as exemplified by GRA80, GRA81 and GRA82 profiles, dynamic release of AP2XI-2 and AP2XII-1 from DNA induced by IAA resulted in a substantial decrease in MORC/HDAC3 enrichment, which enhanced local chromatin hyperaccessibility and led to a concomitant increase in mRNA abundance of target genes (Fig. 6f; Extended Data Fig. 6c). After the addition of IAA, these sites became accessible as they exhibited stronger ATAC-seq signals, presumably a consequence of MORC and HDAC3 eviction from chromatin. This regulatory pattern applied to all families of canonical merozoite genes, regardless of whether they encode proteins destined for organelles or the surface of the parasite (Extended Data Fig. 6c-e). We also observed that this form of control occurred simultaneously for both genes when they were close to each other and arranged head to head (Fig. 6f). Our findings corroborate the long-held hypothesis that AP2 transcription factors in Apicomplexa function as molecular tethers that facilitate the recruitment of chromatin modifiers to specific DNA sequences, which in turn regulates the degree of transcriptional permissiveness of the parasite genome^9, 10, 33^.

## Co-depletion of AP2XI-2 and AP2XII-1 induces a downstream network of secondary transcriptional regulators to guide merogony

In our results, we also identified some genes that exhibited high RNA levels and hyperaccessible chromatin signatures after the addition of IAA, but that lacked the characteristic recruitment of MORC and HDAC3 to their promoters in the untreated state (e.g., PNP, Extended Data Fig. 7a). Conversely, this observation also applies to clusters of tachyzoite genes that are repressed by the simultaneous depletion of AP2XI-2 and AP2XII-1 (Clusters V and VI, Fig. 1f). For those genes, a strong decrease in ATAC-seq signals was observed after the addition of IAA, but no trace of any component of the MORC repressive complex was detected by ChIP in their environment (Fig. 6f; Extended Data Fig. 7b-d). This suggests that AP2XI-2- and AP2XII-1 restrict chromatin accessibility in gene regulatory regions via an indirect mechanism that is not dependent on their DNA-binding activities or their functional partners MORC and HDAC3. This transcriptional output may originate from secondary transcription factors, which in turn dictate the setting of a particular predetermined transcriptional program for a particular stage^1, 10^. This hypothesis is supported by the observation that simultaneous depletion of AP2XI-2 and AP2XII-1 triggers the transcription of seven AP2 and one C2H2 zinc finger TFs, all of which are controlled by the chromatin occupancy dynamics of MORC and HDAC3 (Fig. 6g; Extended Data Fig. 8). A relevant example is AP2IX-1, whose expression is restricted to merozoites and which, when transiently expressed in tachyzoites, has been shown to suppress the expression of SRS29B, which encodes SAG1, the major surface antigen of tachyzoites^36^. Thus, AP2IX-1 likely acts downstream of AP2XI-2 and AP2XII-1 (Extended Data Fig. 8e) and, when expressed, contributes to the restructuring of the SRS repertoire during merogony. Although less frequent in our study, this second wave of TFs may also operate as transcriptional activators to take over the expression of pre-sexual genes not directly regulated by AP2XI-2 and AP2XII-1 (e.g., the *PNP* locus; Extended Data Fig. 7a), indicating that the development to the merozoite stage is subject to an intricate regulatory cascade.

## Discussion

Simultaneous depletion of AP2XII-1 and AP2XI-2 is sufficient to initiate the pre-sexual transcriptional program and silence the tachyzoite determinants in a remarkably more effective manner than depletion of MORC or inhibition of HDAC3. We were able to produce a large number of meronts at all developmental stages in cultured cells. Typically, achieving this level of output would require infecting a significant number of kittens, which poses significant ethical and technical challenges. This approach allowed us to gain a deeper understanding of the biology of pre-gametes, an area that has been largely overlooked so far, and to expand upon what has been inferred from studying other parasites of the same phylum. We provided compelling evidence that *Toxoplasma* schizonts produced *in vitro*, undergo endopolygeny with karyokinesis during merogony. Compared to other Coccidian parasites phylogenetically related to *Toxoplasma*, this process of division is similar to the porcine parasite *Cystoisospora suis* but differs from the endopolygeny process without nuclear fission observed in *S. neurona*^30, 37^. All predefined morphotypes (A to E) have been detected over time *in vitro*, including type E meronts, which are expected to mature into sexual gametes^3^. However, fully differentiated microgametocytes and macrogametocytes were not observed probably because gamete formation requires a complex genetic program as in *Plasmodium falciparum*^38, 39^ or a dedicated metabolic environment found exclusively in feline enterocytes^40^.

Merozoites differ markedly from tachyzoites in that they express a much smaller but specific repertoire of MIC, ROP, and GRA proteins. It is noteworthy that the pregametes have a more limited host cell range for infection as opposed to tachyzoites, which can invade and develop in almost any nucleated cells of a warm-blooded host. Accordingly, merozoites that are fully differentiated *in vitro* are unable to egress, and even when prompted to do so, their ability to glide and infect human fibroblasts is impaired (Extended Data Fig. 2f). This suggests that these *in vitro* differentiated merozoites may have a similar migratory ability as those found in infected cat gut and can move through the gastrointestinal tract and within mucus layers, however, they lack the capability to invade and spread on fibroblast surfaces as tachyzoites do^12^. This finding aligns with the remodeling of their surface proteins (e.g., SRS and family A), which have been proposed to play an important role in gamete recognition and fertilization, similar to *Plasmodium*’s 6-cys protein family^41^.

Mechanistically, AP2XII-1 and AP2XI-2 bind cooperatively to DNA as homo- and heterodimers and selectively recruit HDAC3 and MORC to the promoter of merozoite genes, which in turn create a non-permissive chromatin environment for transcription in tachyzoites. In this process, MORC likely forms dimers that topologically entrap DNA loops^42^ and support chromatin condensation through synergistic covalent modification of nucleosomes catalyzed by its partner, the histone deacetylase HDAC3^1^. Our model was supported by a recent report showing how MORC proteins condense chromatin, reduce DNA accessibility to TFs, and thereby repress gene expression in the plant *A. thaliana*^43^. AP2XII-1 and AP2XI-2 co-depletion also drives hierarchical expression of secondary AP2s, all of which have restricted expression at pre-sexual stages and likely act as downstream activators or repressors during merogony. In this hierarchy, AP2IV-3 is singularly conserved in the phylum (Extended Data Fig. 8b), as it shares a homologous DNA-binding domain with AP2-G, the major transcriptional regulator of gametocytogenesis in *P. falciparum*, underscoring a possible convergence of functions within Apicomplexa. This division of responsibilities between primary and secondary TFs results in a tailored transcriptional response that promotes the unidirectional nature of the life cycle. We anticipate that additional AP2s, which may or may not be associated with MORC, are operating downstream in the parasite life cycle, for example in controlling sex determination. Fine-tuning the activity of these regulators in mature merozoites grown *in vitro* could pave the way for the production of functional gametes, opening the doors to *in vitro* fertilization and forward genetics.

## Methods

### Parasites and human cell cultures

Human primary fibroblasts (HFFs, ATCC® CCL-171™) were cultured in Dulbecco’s Modified Eagle Medium (DMEM) (Invitrogen) supplemented with 10% heat-inactivated fetal bovine serum (FBS) (Invitrogen), 10 mM (4-(2-hydroxyethyl)-1-piperazine-ethanesulfonic acid) (HEPES) buffer pH 7.2, 2 mM L-glutamine and 50 μg/ml penicillin and streptomycin (Invitrogen). Cells were incubated at 37°C and 5% CO2. *Toxoplasma* strains used in this study and listed in Supplementary Table 1 were maintained *in vitro* by serial passage on monolayers of HFFs. Cultures were free of mycoplasma as determined by qualitative PCR.

### Reagents

The following primary antibodies were used in the immunofluorescence, immunoblotting, and/or ChIP assays: rabbit anti-TgHDAC3 (RRID: AB_2713903), rabbit anti-TgGAP45 (gift from Pr. Dominique Soldati), mouse anti-HA tag (Roche, RRID: AB_2314622), rabbit anti-HA Tag (Cell Signaling Technology, RRID: AB_1549585), rabbit anti-mCherry (Cell Signaling Technology, RRID: AB_2799246), rabbit anti-FLAG (Cell Signaling Technology, RRID: AB_2798687), mouse anti-MYC clone 9B11 (RRID: AB_2148465), H3K9me3 (Diagenode, RRID: AB_2616044), rabbit Anti-acetyl-Histone H4, pan (Lys 5,8,12) (Millipore, RRID:AB_310270), rat anti-IMC7 (gift from Pr. Gubbels MJ), mouse anti-IMC1 (gift from Pr. Ward GE), mouse anti-AtRx antibody clone 11G8^26^, mouse anti-GRA11b^14^. We have also raised homemade antibodies against linear peptides in rabbits corresponding to the following proteins: MORC_Peptide2 (C+SGAPIWTGERGSGA); AP2XI-2 (C+HAFKTRRTEAAT) TGME49_273980/GRA80 (C+RPPWAPGAGPEN); TGME49_243940/GRA81 (C+QKELAEVAQRALEN); TGME49_277230/GRA82 (C+SDVNTEGDATVANPE); TGME49_209985/ROP26 (CQETVQGNGETQL); SRS48 family (CKALIEVKGVPK); SRS59B/K (C+IHVPGTDSTSSGPGS); TGME49_314250/BRP1 (C+QVKEGTKNNKGLSDK); TGME49_307640/CK2 kinase (C+IRAQYHAYKGKYSHA); and TGME49_306455 (C+DGRTPVDRVFEE). They were manufactured by Eurogentec and used for immunofluorescence, immunoblotting and/or chromatin immunoprecipitation. Secondary immunofluorescent antibodies were coupled with Alexa Fluor 488 or Alexa Fluor 594 (Thermo Fisher Scientific). Secondary antibodies used in Western blotting were conjugated to alkaline phosphatase (Promega) or horseradish peroxidase.

### Auxin-induced degradation

Degradation of AP2XII-1-AID-HA, AP2XI-2-mAID-HA, and AP2XII-1-AID-HA /AP2XI-2-mAID-MYC was achieved with 3-indoleacetic acid (IAA, Sigma-Aldrich # 45533). A stock of 500 mM IAA dissolved in 100% EtOH at a ratio of 1:1,000 was used to degrade mAID-tagged proteins to a final concentration of 500 μM. The mock treatment consisted of an equivalent volume of 100% EtOH at a final concentration of 0.0789% (wt/vol). To monitor the degradation of AID-tagged proteins, parasites grown in HFF monolayers were treated with auxin or ethanol alone for various time intervals at 37 °C. After treatment, parasites were harvested and analyzed by immunofluorescence or Western blotting.

### Immunofluorescence microscopy

*Toxoplasma*-infected HFF cells grown on coverslips were fixed in 3% formaldehyde for 20 minutes at room temperature, permeabilized with 0.1% (v/v) Triton X-100 for 15 minutes, and blocked in phosphate buffered saline (PBS) containing 3% (w/v) BSA. Cells were then incubated with primary antibodies for 1 hour, followed by the addition of secondary antibodies conjugated to Alexa Fluor 488 or 594 (Molecular Probes). Nuclei were stained with Hoechst 33258 (2 μg/ml in PBS) for 10 minutes at room temperature. After washing four times in PBS, coverslips were mounted on a glass slide with Mowiol mounting medium, and images were acquired with a fluorescence microscope ZEISS ApoTome.2 and processed with ZEN software (Carl Zeiss, Inc.).

For IFA of *in vivo* cat stages, small intestines of infected kittens from a previous study^15^ embedded in paraffin were sectioned to 3 μm and dried over night at 37 °C. Deparaffinization was performed first 3 times 2 min in xylene, washed twice for 1 min in 100% ethanol and finally rehydrated sequentially 1 min in 96% and 70% ethanol and water. For antigen retrieval, samples were boiled in a pressure cooker for 20 min in citrate buffer pH 6.1 (Dako Target Retrieval Solution, S2369) and transferred to water. Cells were permeabilized in 0.3%Triton X-100/PBS and blocked with FCS. Staining was performed over night at 4 °C using either mouse anti-GRA11b^14^ and rabbit anti-IMC1 (gift from Pr. Dominique Soldati) or rabbit anti-GRA80 with rat immune serum in 20% FCS/0.3% TritonX-100/PBS. The samples were then washed and incubated with 1 μg/ml DAPI/20% FCS/0.3% TritonX-100/PBS and either anti-rabbit Alexa Fluor 488 (Invitrogen, A11070) and anti-mouse Alexa Fluor 594 (Invitrogen, A11005) or anti-rat Alexa Fluor 488 (Invitrogen, A11006) with anti-rabbit Alexa Fluor 594 (Invitrogen, A11072) for 1 h at room temperature. After three washes, samples were mounted with Vectashield and imaged either with a Leica DMI 6000 B epi-fluorescent microscope or a Leica SP8 confocal microscope. Confocal images were deconvoluted using SVI Huygens Professional. Maximum intensity projections were performed using FIJI 2.9.1.

### Transmission electron microscopy

For ultrastructural observations, *Toxoplasma*-infected HFF grown as monolayers on a 6-well dish were exposed to 500 µM IAA or ethanol solvent as described above before fixation 24 hours or 40 hours post-infection in 2.5% glutaraldehyde in 0.1 mM sodium cacodylate (pH7.4) and processed as described previously^44^. Ultrathin sections of infected cells were stained with osmium tetraoxide before examination with Hitachi 7600 EM under 80 kV equipped with a dual AMT CCD camera system.

### Western blot

Immunoblot analysis of protein was performed as described in Swale *et al.* (2022)^45^. Briefly, ∼10^7^ cells were lysed and sonicated in 50 μl lysis buffer (10 mM Tris-HCl, pH6.8, 0.5% SDS [v/v], 10% glycerol [v/v], 1 mM EDTA, and protease inhibitor cocktail). Proteins were separated using SDS-PAGE, transferred by liquid transfer to a polyvinylidene fluoride membrane (Immobilon-P; EMD Millipore), and Western blots probed with the appropriate primary antibodies and alkaline phosphatase- or horseradish peroxidase-conjugated secondary goat antibodies. Signals were detected using NBT-BCIP (Amresco) or an enhanced chemiluminescence system (Thermo Scientific).

### Plasmid construction

The plasmids and primers used in this work for the gene of interest (GOI) are listed in Supplementary Table 1. To construct the vector pLIC-GOI-HAFlag and pLIC-GOI-mAID-HA or pLIC-GOI-mAID-(MYC)2, the coding sequence of GOI was amplified with primers LIC-GOI-Fwd and LIC-GOI-Rev using genomic Toxoplasma DNA as template. The resulting PCR product was cloned into the vectors pLIC-HF-dhfr or pLIC-mCherry-dhfr using the ligation-independent cloning (LIC) method. Specific gRNA for GOI, based on the CRISPR/cas9 editing method, was cloned into plasmid pTOXO_Cas9-CRISPR**^1^**. Twenty mers oligonucleotides corresponding to specific GOI were cloned using the Golden Gate strategy. Briefly, the primers GOI-gRNA-Fwd and GOI-gRNA-Rev containing the sgRNA targeting the genomic sequence GOI were phosphorylated, annealed, and ligated into the pTOXO_Cas9-CRISPR plasmid linearized with BsaI, resulting in pTOXO_Cas9-CRISPR::sgGOI.

### *Toxoplasma* transfection

Parasite strains were electroporated with vectors in Cytomix buffer (120 mM KCl, 0.15 mM CaCl2, 10 mM K2HPO4/ KH2PO4 pH 7.6, 25 mM HEPES pH7.6, 2 mM EGTA, 5 mM MgCl2) using a BTX ECM 630 machine (Harvard Apparatus). Electroporation was performed in a 2 mm cuvette at 1,100 V, 25 Ω, and 25 µF. Antibiotics (concentration) used were chloramphenicol (20 µM), mycophenolic acid (25 µg/ml) with xanthine (50 µg/ml), pyrimethamine (3 µM), or 5-fluorodeoxyuracil (10 µM) as needed. Stable transgenic parasites were selected with the appropriate antibiotic, cloned in 96-well plates by limiting dilution, and verified by immunofluorescence assay or genomic analysis.

### Chromatographic purification of FLAG tagged proteins

*Toxoplasma* extracts from RHΔku80 or PruΔku80 cells stably expressing HAFlag-tagged AP2XII-1 or AP2XI-2 proteins, respectively, were incubated with anti-FLAG M2 affinity gel (Sigma-Aldrich) for 1 hour at 4°C. Beads were washed with 10-column volumes of BC500 buffer (20 mM Tris-HCl, pH 8.0, 500 mM KCl, 20% glycerol, 1 mM EDTA, 1 mM DTT, 0.5% NP-40, and protease inhibitors). Bound polypeptides were eluted stepwise with 250 μg/ml FLAG peptide (Sigma Aldrich) diluted in BC100 buffer. For size-exclusion chromatography, protein eluates were loaded onto a Superose 6 HR 10/30 column equilibrated with BC500. The flow rate was set at 0.35 ml/min, and 0.5-ml fractions were collected.

### MS-based proteomic analyses of interactomes and SEC fractions

Protein bands were excised from colloidal blue stained gels (Thermo Fisher Scientific) before in-gel digestion using modified trypsin (Promega, sequencing grade) as previously described**^1^**. Resulting peptides were analyzed by online nanoliquid chromatography (UltiMate 3000 RSLCnano, Thermo Scientific) coupled to tandem MS (Q-Exactive Plus, Q-Exactive HF and Orbitrap Exploris 480, Thermo Scientific, for respectively AP2XI-2 interactome, AP2XII-1 interactome, and SEC fractions). Peptides were sampled on a 300 µm x 5 mm PepMap C18 precolumn and separated on a 75 µm x 250 mm C18 column (Reprosil-Pur 120 C18-AQ, 1.9 μm, Dr. Maisch, for AP2XI-2 interactome, and Aurora, 1.7 µm, IonOpticks, for AP2XII-1 interactome and SEC fractions) using 25-min gradients. MS and MS/MS data were acquired using Xcalibur version 4.0 (Thermo Scientific). Peptides and proteins were identified using Mascot (version 2.8.0) through concomitant searches against the *Toxoplasma gondii* database (ME49 taxonomy, version 58 downloaded from ToxoDB), the Uniprot database (*Homo sapiens* taxonomy for interactomes or *Trichoplusia ni* for SEC fractions, 20220527 download), and a homemade database containing the sequences of classical contaminant proteins found in proteomic analyses (human keratins, trypsin…). Trypsin/P was chosen as the enzyme and two missed cleavages were allowed. Precursor and fragment mass error tolerances were set at respectively at 10 and 20 ppm. Peptide modifications allowed during the search were: Carbamidomethyl (C, fixed), Acetyl (Protein N-term, variable) and Oxidation (M, variable). The Proline software (version 2.2.0) was used for the compilation, grouping, and filtering of the results (conservation of rank 1 peptides, peptide length ≥ 6 amino acids, false discovery rate of peptide-spectrum-match identifications < 1%, and minimum of one specific peptide per identified protein group). Intensity-based absolute quantification (iBAQ) values were calculated for each protein group in Proline using MS1 intensities of specific and razor peptides.

### MS-based quantitative analyses of parasite proteomes

HFF cells were grown to confluence, infected with RH AP2XII-1 KD / AP2XI-2 KD strain and treated with IAA for 24h, 32h and 48h or mock-treated. Three biological replicates were prepared and analyzed for each condition. Proteins were extracted using the Cell lysis buffer (Invitrogen). Seven micrograms of proteins were then stacked in the top of a 4-12% NuPAGE gel (Invitrogen), stained with Coomassie blue R-250 (Bio-Rad) before in-gel digestion using modified trypsin (Promega, sequencing grade) as previously described**^1^**. The resulting peptides were analyzed by online nanoliquid chromatography coupled to MS/MS (Ultimate 3000 RSLCnano and Q-Exactive HF, Thermo Fisher Scientific) using a 360-min gradient. For this, peptides were sampled on a 300 μm × 5 mm PepMap C18 precolumn and separated in a 200 cm µPAC column (PharmaFluidics). MS and MS/MS data were acquired using the Xcalibur software version 4.0 (Thermo Scientific). Peptides and proteins were identified by Mascot (version 2.8.0, Matrix Science) through concomitant searches against the *Toxoplasma gondii* database (ME49 taxonomy, version 58 downloaded from ToxoDB), the Uniprot database (*Homo sapiens* taxonomy, 20220527 download), and a homemade database containing the sequences of classical contaminant proteins found in proteomic analyses (human keratins, trypsin…). Trypsin/P was chosen as the enzyme and two missed cleavages were allowed. Precursor and fragment mass error tolerances were set at respectively at 10 and 20 ppm. Peptide modifications allowed during the search were: Carbamidomethyl (C, fixed), Acetyl (Protein N-term, variable) and Oxidation (M, variable). The Proline software (version 2.2.0) was used for the compilation, grouping, and filtering of the results (conservation of rank 1 peptides, peptide length ≥ 6 amino acids, false discovery rate of peptide-spectrum-match identifications < 1%, and minimum of one specific peptide per identified protein group). Proline was then used to perform a MS1 label-free quantification of the identified protein groups based on razor and specific peptides. Statistical analyses were performed using ProStaR^49^. Peptides and proteins identified in the reverse and contaminant databases or matched to human sequences were discarded. Only proteins identified by MS/MS in a minimum of two replicates of one condition and quantified in the three replicates of one condition were conserved. After log2 transformation, abundance values were normalized using the variance stabilizing normalization (vsn) method, before missing value imputation (SLSA algorithm for partially observed values in the condition and DetQuantile algorithm for totally absent values in the condition). For comparison of each IAA-treated conditions to mock-treated condition, statistical testing was conducted with limma, whereby differentially expressed proteins were selected using a log2(Fold Change) cut-off of 1 and a p-value cut-off of 0.01, allowing to reach a false discovery rate inferior to 5% according to the Benjamini-Hochberg estimator. Proteins found differentially abundant but identified by MS/MS in less than two replicates, and detected in less than three replicates, in the condition in which they were found to be more abundant were invalidated (p-value = 1). Protein abundances measured in the four different conditions were also compared globally by analysis of variance (ANOVA) using Perseus; q-values were obtained by Benjamini-Hochberg correction.

### Chromatin immunoprecipitation coupled with ilumina sequencing

#### Chromatin immunoprecipitation

HFF cells were grown to confluence and infected with KD strains as indicated in the figure legends. Harvested intracellular parasites were cross-linked with formaldehyde (final concentration 1%) for 8 minutes at room temperature, and cross-linking was stopped by addition of glycine (final concentration 0.125 M) for 5 minutes at room temperature. The parasites were lysed in ice-cold lysis buffer A (50 mM HEPES KOH pH7.5, 14 0mM NaCl, 1 mM EDTA, 10% glycerol, 0.5% NP-40, 0.125% Triton X-100, protease inhibitor cocktail) and after centrifugation, cross-linked chromatin was sheared in buffer B (1 mM EDTA pH 8.0, 0.5 mM EGTA pH 8.0, 10 mM Tris pH 8.0, protease inhibitor cocktail) by sonication with a Diagenode Biorupter. Samples were sonicated for 16 cycles (30 seconds ON and 30 seconds OFF) to achieve an average size of 200-500 base pairs. Sheared chromatin, 5% BSA, a protease inhibitor cocktail, 10% Triton X-100, 10% deoxycholate, magnetic beads coated with DiaMag protein A (Diagenode), and antibodies against epitope tags (HA or MYC) or the protein of interest (MORC or HDAC3) were used for immunoprecipitation. A rabbit IgG antiserum served as a control mock. After overnight incubation at 4°C on a rotating wheel, chromatin-antibody complexes were washed and eluted from the beads using the iDeal ChIP-seq kit for transcription factors (Diagenode) according to the manufacturer’s protocol. Samples were de-crosslinked by heating for 4 hours at 65 °C. DNA was purified using the IPure kit (Diagenode) and quantified using Qubit Assays (Thermo Fisher Scientific) according to the manufacturer’s protocol. For ChIP-seq, the purified DNA was used for library preparation and subsequently sequenced by Arraystar Co (USA).

#### Library Preparation, Sequencing and Data analysis (Arraystar)

ChIP sequencing libraries were prepared according to the Illumina protocol Preparing Samples for ChIP Sequencing of DNA. Library preparation: 10 ng of DNA from each sample was converted to blunt-end phosphorylated DNA fragments using T4 DNA polymerase, Klenow polymerase, and T4 polymerase (NEB); an ‘A’ base was added to the 3’ end of the blunt-end phosphorylated DNA fragments using the polymerase activity of Klenow (Exo-Minus) polymerase (NEB); Illumina genomic adapters were ligated to the A-tailed DNA fragments; PCR amplification to enrich the ligated fragments was performed using Phusion High Fidelity PCR Master Mix with HF Buffer (Finnzymes Oy). The enriched product of ∼200-700 bp was excised from the gel and purified. Sequencing: the library was denatured with 0.1 M NaOH to generate single-stranded DNA molecules and loaded into flow cell channels at a concentration of 8 pM and amplified in situ using TruSeq Rapid SR cluster kit (# GD-402-4001, Illumina). Sequencing was performed at 100 cycles on the Illumina HiSeq 4000 according to the manufacturer’s instructions.

#### Data Analysis

After the sequencing platform generated the sequencing images, the stages of image analysis and base calling were performed using Off-Line Basecaller software (OLB V1.8). After passing Solexa CHASTITY quality filter, the clean reads were aligned to T. gondii reference genome (TGME49) using BOWTIE V2 then converted and sorted using Bamtools. Aligned reads were used for peak calling of the ChIP enriched peaks using MACS V2.2 with a cutoff p-value of 10^-4^. Data visualization: For IGB visualization and gene centered analysis using Deeptools, MACS2 generated bedgraph files were processed with the following command: “sort-k1,1-k2,2n 5_treat_pileup.bdg > 5_treat_pileup-sorted.bdg” then converted using the BedGraphToBigWig program (ENCODE project). The Deeptools analysis were generated using “computeMatrix reference point” with the following parameters (-- minThreshold 2, --binSize 10 and –averageTypeBins sum). Plotprofile or heatmap was then used with a k-mean clustering when applicable. Inter sample comparison were obtained using the nf-core chip-seq workflow with standard parameters^46^. From this pipeline, HOMER (annotatePeaks) was used to analyze peak distribution relative to gene features. All these raw and processed files can be found at Series GSE222819.

### RNA-seq and sequence alignment

Total RNAs were extracted and purified using TRIzol (Invitrogen, Carlsbad, CA, USA) and RNeasy Plus Mini Kit (Qiagen). RNA quantity and quality were measured by NanoDrop 2000 (Thermo Scientific). For each condition, RNAs were prepared from three biological replicates. RNA integrity was assessed by standard non-denaturing 1.2% TBE agarose gel electrophoresis. RNA sequencing was performed following standard Illumina protocols, by Novogene (Cambridge, United Kingdom). Briefly, RNA quantity, integrity, and purity were determined using the Agilent 5400 Fragment Analyzer System (Agilent Technologies, Palo Alto, California, USA). The RQN ranged from 7.8 to 10 for all samples, which was considered sufficient. Messenger RNAs (mRNA) were purified from total RNA using poly-T oligo-attached magnetic beads. After fragmentation, the first strand cDNA was synthesized using random hexamer primers. Then the second strand cDNA was synthesized using dUTP, instead of dTTP. The directional library was ready after end repair, A-tailing, adapter ligation, size selection, USER enzyme digestion, amplification, and purification. The library was checked with Qubit and real-time PCR for quantification and bioanalyzer for size distribution detection. Quantified libraries will be pooled and sequenced on Illumina platforms, according to effective library concentration and data amount. The samples were sequenced on the Illumina NovaSeq platform (2 x 150 bp, strand-specific sequencing) and generated ∼40 million paired-end reads for each sample. The quality of the raw sequencing reads was assessed using FastQC (www.bioinformatics. babraham.ac.uk/projects/fastqc/) and MultiQC. For the expression data quantification and normalization, the FASTQ reads were aligned to the ToxoDB-49 build of the T. gondii ME49 genome using Subread version 2.0.1 with the following options ‘subread-align -d 50 -D 600 -- sortReadsByCoordinates’. Read counts for each gene were calculated using featureCounts from the Subread package. Differential expression analysis was conducted using DESeq2 and default settings within the iDEP.96 web interface^47^. Transcripts were quantified and normalized using TPMCalculator. The Illumina RNA-seq dataset generated during this study is available at the National Center for Biotechnology Information (NCBI): BioProject # PRJNA921935.

### Nanopore Direct RNA Sequencing (DRS)

The mRNA library preparation followed the SQK-RNA002 kit (Oxford Nanopore)–recommended protocol, the only modification was the input mRNA quantity increased from 500 to 1000 ng, and all other consumables and parameters were standard. Final yields were evaluated using the Qubit HS dsDNA kit (Thermo Fisher Scientific, Q32851) with minimum RNA preps reaching at least 200 ng. For all conditions, sequencing was performed on FLO-MIN106 flow cells either using a MinION MK1C or MinION sequencer. All datasets were subsequently basecalled (high accuracy) with a Guppy version higher than 5.0.1 with a *Q* score cutoff of >7. Long read alignment was performed by Minimap2 as previously described^48^. Sam files were converted to bam and sorted using Samtools 1.4. Alignments were converted and sorted using Samtools 1.4.1. For the three described samples, Toxoplasma aligned reads range between 600,000 and 800,000. The Nanopore DRS dataset is available at the National Center for Biotechnology Information (NCBI): BioProject # PRJNA921935.

### Assay for Transposase-Accessible Chromatin with high-throughput sequencing (ATAC-seq)

Intracellular tachyzoites (non-treated or IAA treated for 24h) were prepared using HFF cells monolayers in a T175 format, which was freshly scrapped, gently homogenised by pipetting and centrifuged at 500*g. Prior to initiating the transposition protocol, the pellet was gently washed with warm dpbs (Life technologies) and resuspended in 500 μl of cold PBS + protease inhibitor (Diagenode kit). Nuclei preparation, permeabilization, Tn5 transposition and library preparation was prepared following precisely the Diagenode ATAC-SEQ kit protocol (C01080002). Nuclei permeabilization was performed on an estimated 100000 tachyzoites by diluting 10 μl of DPBS cell suspension (from one T175 resuspended in 500 μl) in 240 of DPSB + protease inhibitor (1/25 dilution). From this dilution, 50 μl were then taken to perform the transposition reaction. Of note, the permeabilization protocol used a 3 minute 0,02% digitonin (Promega) exposure. Following the Tn5 reaction, libraries were amplified using the Diagenode 24 UDI kit 1 (ref 01011034) following standard protocol procedures. Libraries were multiplexed and sequenced on a single Novaseq6000 lane by Fasteris (Genesupport SA) using 2*50 cycles, generating on average 27 million reads. Following the demultiplexing of raw reads by bcl2fastq V3, trimming, quality control, alignment to the ME49 reference genome (using bwa2) and duplicate read merging (using Picard) was performed by the nf-core ATAQ-SEQ pipeline^46^. Data visualization: For IGB visualization and gene centered analysis using Deeptools, Picard merged bam files were converted to bigWig file format using a bin size of 5 by bamCoverage (Deeptools). The Deeptools analysis were then generated using “computeMatrix reference point” with the following parameters (--minThreshold 2, --binSize 10 and –averageTypeBins sum). Quantitative analysis of UT vs IAA 24h conditions was performed by nf-core through a broad peak calling/annotation (MACS2) followed by HOMER (annotatePeaks) to analyse peak distribution relative to gene features. Reads were counted on annotated peaks by featureCounts and counts were processed by DeSeq2 to generate global statistical analysis of peak intensities in between conditions using biological duplicates. All these raw and processed files can be found at Series GSE222832.

### Gene synthesis for recombinant co-expression of *Tg*AP2XI-2 and *Tg*Ap2XII-I

Gene synthesis for all insect cell codon-optimized constructs was provided by GenScript. Both AP2 genes were cloned within the co-expression donor vector pFastBac dual which accepts two constructs. The AP2XI-2 expression cassette was derived from the TGME49_310900 gene with a fused dual Strep Tag and Tobacco Etch Virus (TEV) site in the N-terminus. AP2XII-1 was derived from the full length TGME49_218960 gene with an additional non-cleavable FLAG-TAG on the C-terminus. The AP2XI-2 expression cassette was under the polyhedrin promoter while AP2XII-1 was under the P10 promoter.

### Generation of baculovirus

Bacmid cloning steps and baculovirus generation were performed using EMBacY baculovirus (gifted by I. Berger), which contains a yellow fluorescent protein reporter gene in the virus backbone. The established standard cloning and transfection protocols set up within the EMBL Grenoble eukaryotic expression facility were used. Although baculovirus synthesis (V0) and amplification (to V1) were performed with SF21 cells cultured in SF900 III medium (Life Technologies), large-scale expression cultures were performed with Hi-5 cells cultured in the protein free ESF 921 Insect Cell Culture Medium (Expression System) and infected with 0.8-1.0% (v/v) of generation 2 (V1) baculovirus suspensions and harvested 72 hours after infection.

### Strep-TEV-AP2XI-2/AP2XII-1-flag expression and purification

For purification, three cell pellets of about 500 ml of Hi-5 culture were each resuspended in 50 ml of lysis buffer [50 mM tris (pH 8.0), 400 mM NaCl, and 2 mM b-mercaptoethanol (BME)] in the presence of an anti-protease cocktail (Complete EDTA-free, Roche) and 1 μl of benzonase (Merck Millipore, 70746). Lysis was performed on ice by sonication for 3 min (30-s on/ 30-s off, 45° amplitude). After the lysis step, 10% of glycerol was added. Clarification was then performed by centrifugation for 1 hour at 12,000*g* and 4°C and vacuum filtration using 45um nylon filter systems (SteriFlip - Merck Millipore). Prior to purification, tetrameric avidin (Biolock - IBA lifescience) was added to the clarified lysate (1/1000 v/v) which was then batch incubated for 20 minutes with 3 ml of Strep-Tactin XT (IBA lifescience). Following the incubation, the resin was retained on a glass column and washed using 10 ml of lysis buffer. The elution was then performed using 1X BXT buffer (IBA lifescience) which contains 50 mM biotin, 100 mM Tris pH 8and 150 mM NaCl. This initial 1X solution was further supplemented with 300 mM NaCL, 2 mM BME and 10 % glycerol. Following Strep-Tactin XT elution, the sample was concentrated to 500 μl using a 100 kDa concentrator (Amicon Ultra 4 - Merck Millipore) injected on an ÄKTA pure FPLC using a Superose 6 increase column 10/300 GL (Cytiva) running in 50 mM Tris pH: 8, 400 mM NaCl, 1 mM BME.

### Software and Statistical analyses

Volcano plots, scatter plots, and histograms were generated with Prism 7. Sample sizes were not predetermined and chosen according to previous literature. Experiments were performed in biological replicates and provided consistent statistically relevant results. No method of randomization was used. All experiments were performed in independent biological replicates as stated for each experiment in the manuscript. All corresponding treatment and control samples from ChIP-seq and RNA-seq were processed at the same time to minimize technical variation. Investigators were not blinded during the experiments.

### Ethics statement

Animal experiments were performed under the direct supervision of a veterinary specialist, and according to Swiss law and guidelines on Animal Welfare and the specific regulations of the Canton of Zurich under permit numbers 130/2012 and 019/2016, as approved by the Veterinary Office and the Ethics Committee of the Canton of Zurich (Kantonales Veterinäramt Zürich, Zollstrasse 20, 8090 Zürich, Switzerland).

## Data availability

Correspondence and requests for materials should be addressed to M.A.H. Nanopore and Illumina RNA Sequencing data that support the findings of this study have been deposited under the BioProject number PRJNA921935. The ChIPseq and ATAC-seq data have been deposited to the GEO Datasets under accession number GSE222819 and GSE222832,respectively. The MS proteomics data have been uploaded to the ProteomeXchange Consortium via the PRIDE partner repository with the dataset identifiers PXD039400 and PXD039390 for respectively the proteome-wide and interactome analyses. Processed proteomics data is available in Supplementary Table 3.

## Acknowledgments

We are grateful to the developers of the ToxoDB.org Genome Resource. ToxoDB and EuPathDB are part of the National Institutes of Health/National Institutes of Allergy and Infectious Diseases (NIH/NIAID)-funded Bioinformatics Resource Center. We thank the excellent technical staff of the Electron Microscopy Core Facility at the Johns Hopkins University School of Medicine Microscopy Facility. This work was supported by the Laboratoire d’Excellence (LabEx) ParaFrap [ANR-11-LABX-0024], the Agence Nationale pour la Recherche [Project HostQuest, ANR-18-CE15-0023], [Project ApiNewDrug, ANR-21-CE35-0010-01], [Project ApiMORCing, ANR-21-CE15-0002-01], and Fondation pour la Recherche Médicale [FRM Equipe, EQU202103012571]. I.C. was supported by a NIH grant [R01 AI060767]. MS-based proteomic experiments were partially supported by Agence Nationale de la Recherche under projects ProFI (Proteomics French Infrastructure, ANR-10-INBS-08) and GRAL, a program from the Chemistry Biology Health (CBH) Graduate School of University Grenoble Alpes (ANR-17-EURE-0003). We express our gratitude to Pr. Gubbels MJ for his generous donation of antibodies, including the rat anti-IMC7.

## Author Contributions

M.-A.H. supervised the research and coordinated the collaboration. A-V.A. M.S., C.S., D.C.F., A.B., D.C., C.C., and M.-A.H. designed, performed and interpreted the experimental work. Specifically, Y.C. and C.B. performed the mass spectrometric analyzes. IC performed and interpreted TEM. C.R. and A.B.H. performed confocal microscopy of infected cat mucosa. M.-A.H., wrote the paper with editorial support from I.C. and comments from all other authors.

