## Supplementary Figures for "*In vitro* production of cat-restricted *Toxoplasma* pre-sexual stages by epigenetic reprogramming"

A total of 4 supplementary figures are included here with legends.

#### Supplementary Tables

A total of 6 tables are submitted as separate SI Excel files and their legend are included here.

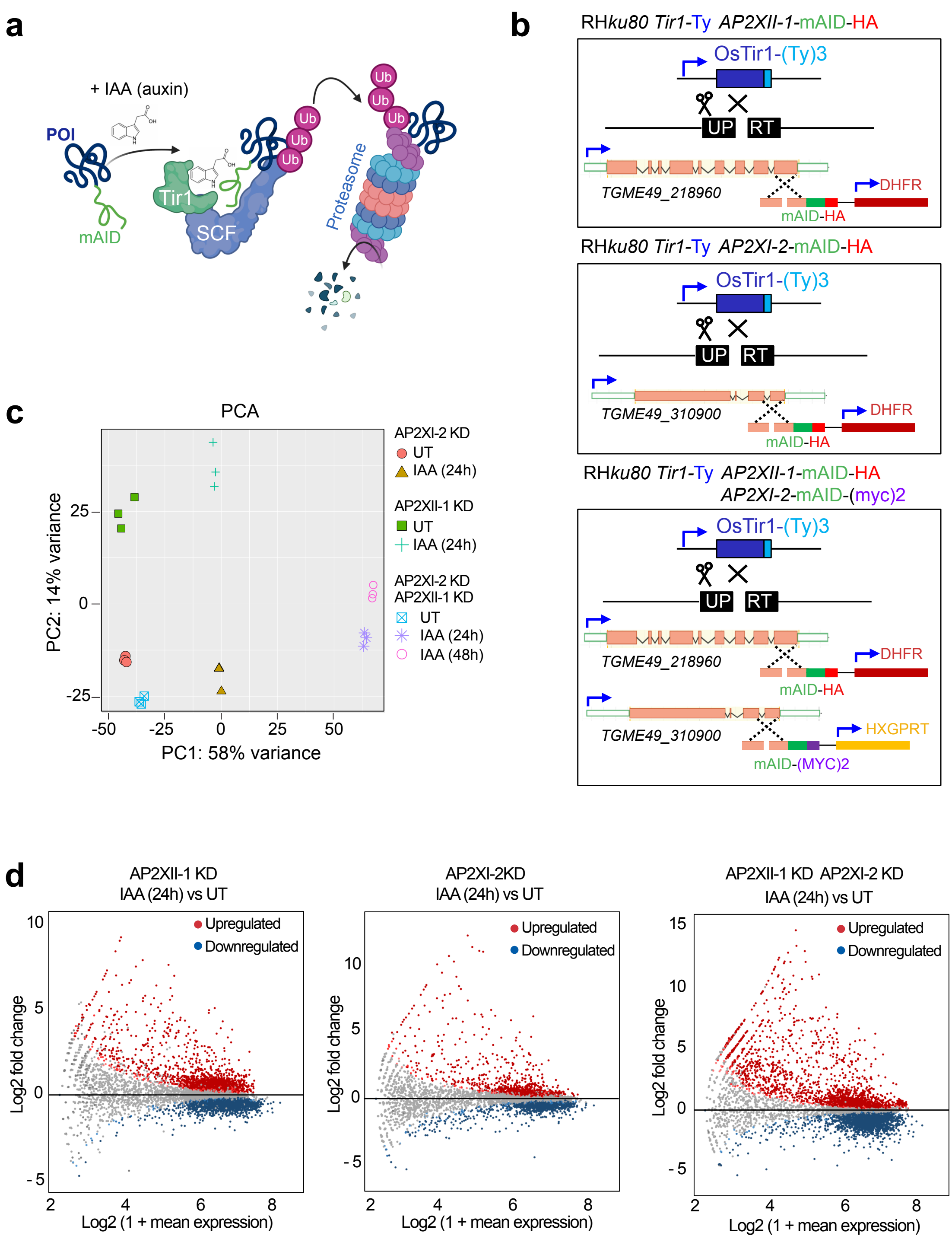


**Supplementary Fig. 1 | Depletion of AP2XII-1 and AP2XI-2 alone or together disrupts expression of a variety of *Toxoplasma* genes.** **a**, Schematic representation of the auxin-induced degron system. The plant auxin receptor, transport inhibitor response 1 (TIR1), is ectopically expressed in *Toxoplasma*. It is then incorporated into the parasitic SCF complex. In the presence of the auxin hormone indole-3-acetic acid (IAA), TIR1 associates with the AID, which is fused to the protein of interest (POI). SCF /TIR1 recruits an E2 ligase and adds a chain of ubiquitin molecules to AID, leading to degradation of POI by the proteasome. **b**, We chose the UPRT locus to integrate TIR1 under the control of a promoter that allows mild expression of the protein tagged to Ty (Farhat et al. 2020). We used a mini AID (mAID) tagging LIC system for conditional AP2XII-1 and AP2XI-2 single depletion or co-depletion. The resulting cell lines are shown. **c**, Principal component analysis (PCA) shows biological and technical variability between samples after Illumina sequencing of mRNA extracted in triplicate from single KD or double KD parasite samples untreated or treated with IAA for 24 or 48 hours. **d**, MA plot of the log2 fold change versus log2 mean expression of all genes showing differential gene expression before and after addition of IAA in three different parasite knockdown lineages.

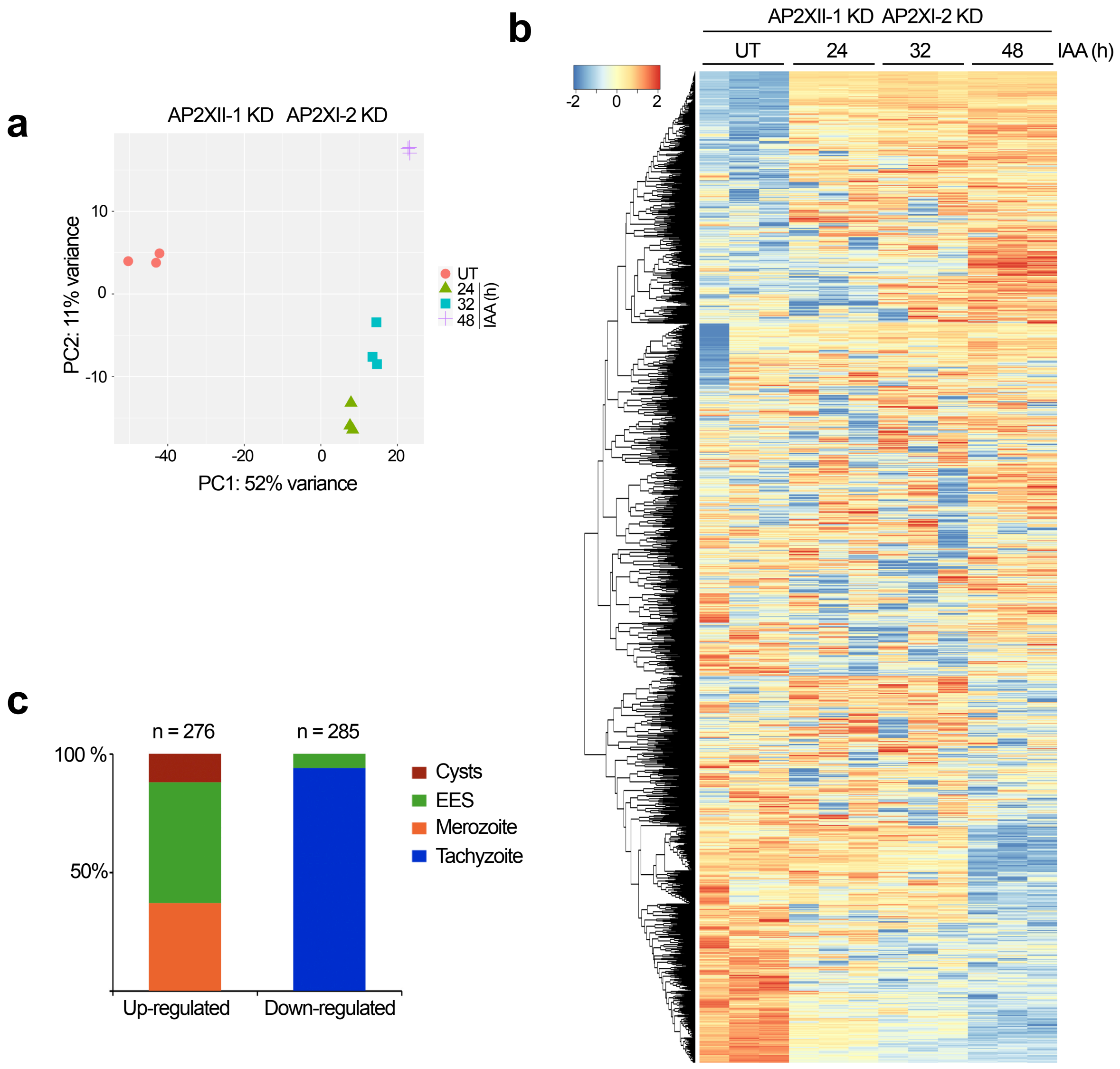


**Supplementary Fig. 2 | The simultaneous depletion of AP2XII-1 and AP2XI-2 disrupts drastically the proteins levels. a,** Principal component analysis (PCA) shows biological and technical variability between proteome samples extracted in triplicate after 24, 32, and 48 hours of AP2XII-1/AP2XI-2 KD induction and compared with the untreated sample (UT). **b**, Hierarchical clustering with iDEP.90 (Ge et al., 2018) of protein profiles of parasites untreated or treated with IAA for 24, 32, and 48 hours, where each column denotes a replicate and each row denotes a protein (all 3020 quantified proteins). **c**, Histogram showing the distribution of up- and down-regulated proteins (n= 276 and 285, respectively) after knockdown of AP2XII-1 and AP2XI-2 according to their life stage affiliation.

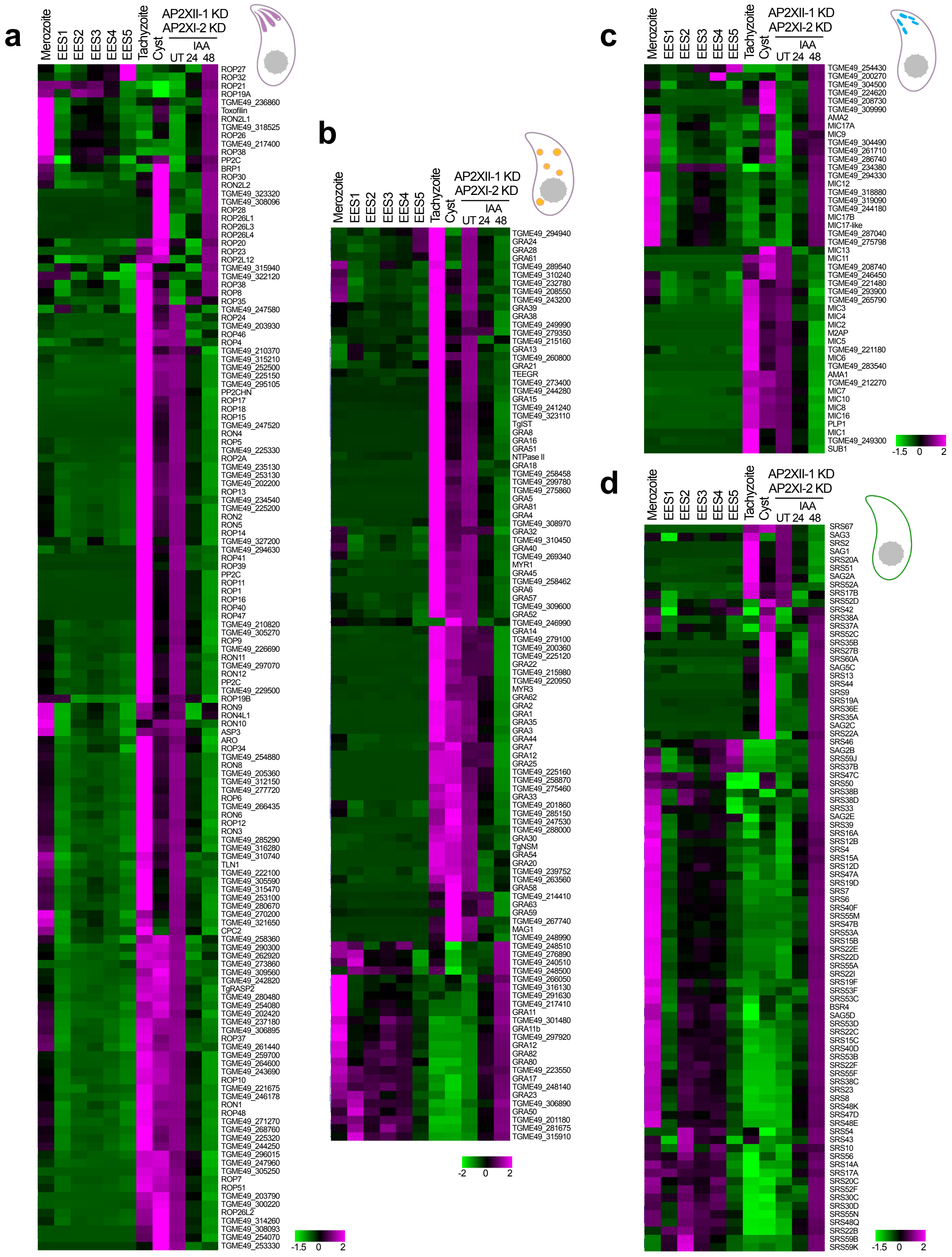


**Supplementary Fig. 3 | Simultaneous AP2XI-2 and AP2XII-1 degradation shifts the transcriptome toward that of merozoites.** Heat map showing hierarchical clustering analysis (Pearson correlation) of selected rhoptry (**a**), dense granule (**b**), microneme (**c**), and SRS (**d**) mRNA transcripts differentially regulated after simultaneous and conditional depletion of AP2XII-1 and AP2XI-2. Shown is the abundance of their transcripts at different developmental stages, namely tachyzoite, cyst, merozoite, and EES. The color scale indicates the log2-transformed fold changes.

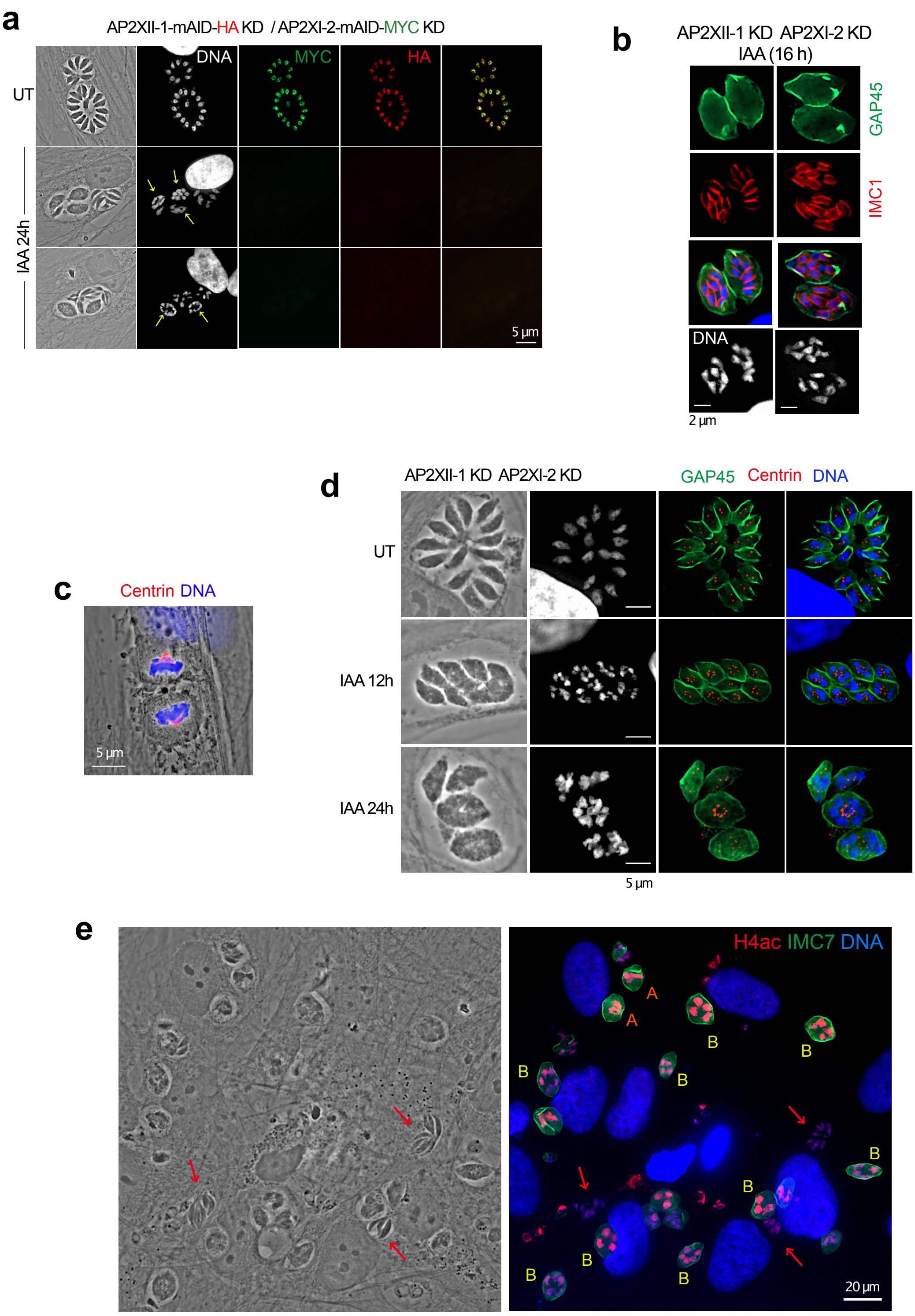


**Supplementary Fig. 4 | Microscopic characterization of endopolygeny after simultaneous degradation of AP2XII-1 and AP2XI-2. a**, IFA of tachyzoites (UT) and AP2XII-1/AP2XI-2-depleted zoites (24 hours post-IAA) were fixed and stained with HA (red) and MYC (green) to detect AP2XII-I-mAID- HA and AP2XI-2-mAID- MYC, respectively. Cells were stained with Hoechst DNA-specific dye (white). Polyploid meronts are indicated by a yellow arrow. **b,** AP2XII-1/AP2XI-2-depleted polyploid meronts (16 hours post-IAA) were fixed and stained with IMC1 (red) and GAP45 (green). **c**, Centrin distribution (red) within the mother fibroblast cell undergoing cytokinesis. **d**, IFA of tachyzoites (UT) and AP2XII-1/AP2XI-2-depleted polyploid meronts (24 hours post-IAA) were fixed and co-stained with centrin (red), GAP45 (green) and Hoechst DNA-specific dye (white or blue). **e,** Close-up of the diversity of pre-gametes stages after co-depletion AP2XII-1 and AP2XI-2 (16 hours post-IAA). Zoite and host cell nuclei are stained with pan-acetylated histone H4 (red) and Hoechst DNA-specific dye (blue), whereas polyploid meronts are labelled with IMC7. Red arrows indicate fully developed merozoites.

### Supplementary Table legends

**Supplementary Table 1 | Description of *T. gondii* Strains, Plasmids, Primers and DNA synthesis.** List of *T. gondii* parasite lines as well as plasmids used in this work. Primers and DNA synthesis construct used in this work are also charted in the table.

**Supplementary Table 2 | AP2XII-1- and APX -2-regulated transcriptomes.** Gene expression profiles in HFF showing that T. gondii genes are differentially regulated by AP2XII-1 and AP2XI-2 using different KD and conditions of IAA induction. RAW and TPM values are given for the indicated samples.

**Supplementary Table 3 | MS-based quantitative analysis of total proteome from T. gondii depleted or not for AP2XII-1 and AP2XI-2.** The proteomes from the AP2XII-1 KD / AP2XI-2 KD strain infecting HFF cells and treated (T) or not (UT) with IAA for 24h, 32h and 48h were analyzed by MS-based label-free quantitative proteomics (three biological replicates per condition). The quantification of proteins (log2 of filtered, normalized and imputed abundances of the different proteins in the different samples are given in columns O to Z) was based on razor and specific peptides (values indicated in column G). Statistical significance was tested using limma for two-by-two sample comparison ; differentially abundant proteins were defined by a log2(fold change) $\geq 1$ or $\leq -1$ and a p-value $\leq 0.01$, allowing to reach a false-discovery rate $< 5$ % according to the Benjamini-Hochberg estimator. A global comparison of the abondances of each protein in the four analyzed conditions was performed using ANOVA (column N).

**Supplementary Table 4 | MS-based characterization of AP2XI-2 interactome.** Total protein extracts from T. gondii cells infecting HFF cells and stably expressing HAFlag-tagged AP2XI-2 protein were submitted to Flag immunoprecipitation and the eluted proteins were separated by SDS-PAGE before staining with Coomassie blue. The protein content of 8 protein bands were submitted to MS-based characterization. The results obtained for each band are presented in the corresponding data sheets.

**Supplementary Table 5 | MS-based characterization of AP2XII-1 interactome.** Total protein extracts from T. gondii cells infecting HFF cells and stably expressing HAFlag-tagged AP2XII-1 protein were submitted to Flag immunoprecipitation and the eluted proteins were separated by SDS-PAGE before staining with Coomassie blue. The protein content of 11 protein bands were submitted to MS-based characterization. The results obtained for each band are presented in the corresponding data sheets.

**Supplementary Table 6 | MS-based characterization of AP2XI-2 interactome using protein expressed in insect cells.** Total protein extracts from insect cells expressing Strep-tagged AP2XI-2 protein were submitted to immunoprecipitation. The eluates were then submitted to size-exclusion chromatography and the proteins in each fraction were separated by SDS-PAGE before staining with Coomassie blue. The protein content in the major protein band of the two main fractions was analyzed by MS-based proteomics. The results obtained for each band are presented in the corresponding data sheets.
