## Extended Data Figures for "*In vitro* production of cat-restricted *Toxoplasma* pre-sexual stages by epigenetic reprogramming"

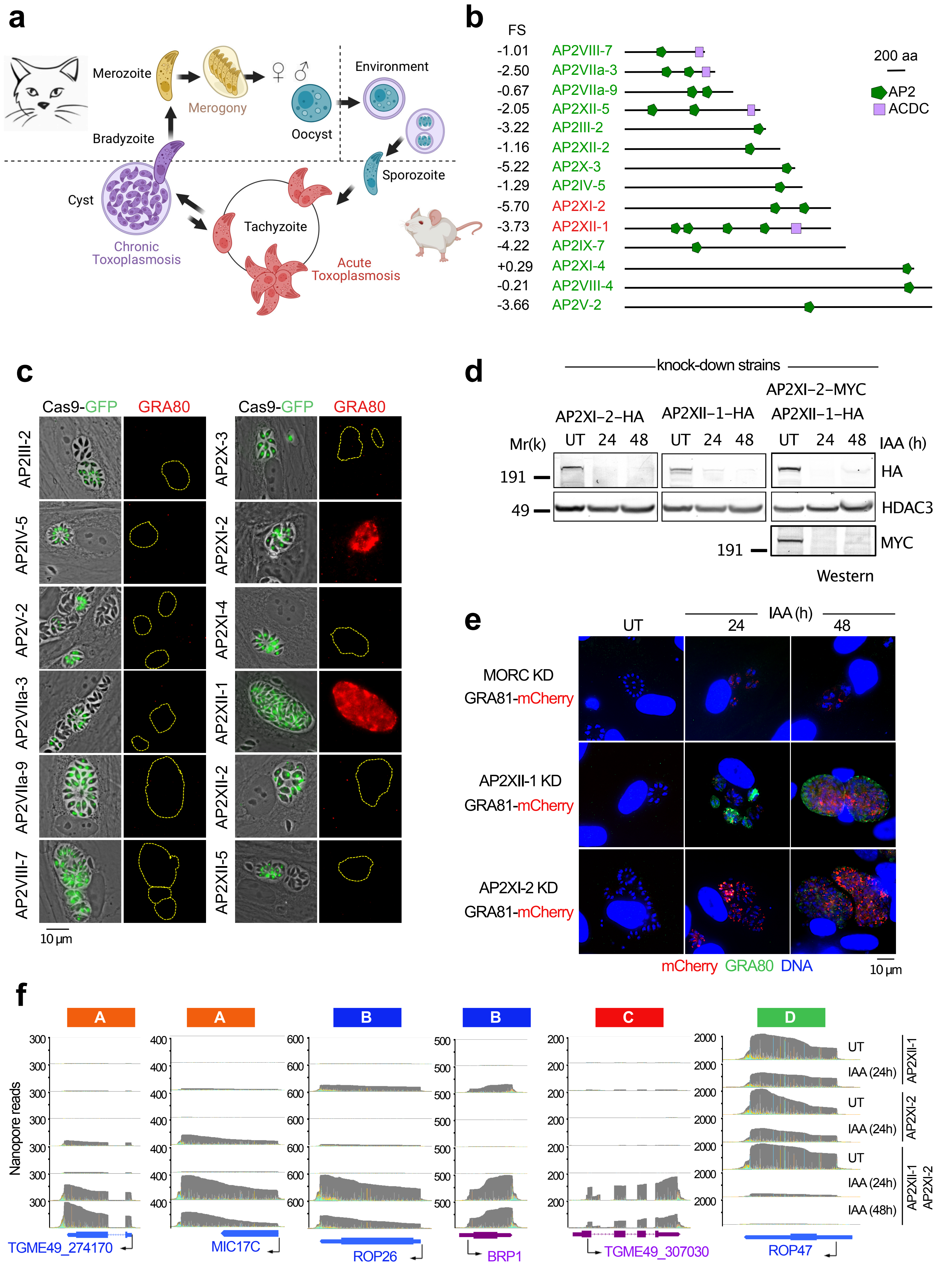


**Extended data Fig. 1**

**Extended Data Fig. 1 | AP2XII-1 alongside AP2XI-2 suppresses the expression of genes specific for the pre-sexual stages.** **a**, Schematic representation of the multistage life cycle of T. gondii. The enteroepithelial cycle begins after ingestion of tissue cysts by felids, with merozoites initiating the formation of gametes after asexual replication cycles, which in turn produce transmissible oocysts that sporulate when exposed to environmental oxygen. After ingestion, the sporozoites are released from the oocysts and differentiate into tachyzoites that cause acute infection. In response to the host immune response, the tachyzoites transform into bradyzoites that form cysts in deep tissues such as the host brain. Predation of a cyst-containing animal by a cat completes the life cycle of the parasite. **b**, Representative domain architectures of MORC partners identified originally by immunoprecipitation coupled to MS-based proteomic analysis. Domains were predicted by SMART and PFAM: AP2 (APETALA2) and ACDC (AP2-Coincident Domain mainly at the Carboxy-terminus). **c**, Representative photomicrographs show intracellular parasites in which AP2 genes were disrupted by transient transfection of the corresponding CRISPR/Cas9 plasmid. Disruption was monitored by Cas9-GFP expression (in green). The level of the merozoite marker GRA80 (in red) was monitored in AP2-deleted zoites (GFP-positive). **d**, Time-course analysis of the expression levels of AP2XII-1 and AP2XI-2 in the single and double KD strains. Samples were collected at the indicated time points after addition of IAA and probed with antibodies against HA, MYC, and HDAC3. The same experiment was repeated three times, and a representative blot is shown. e, IFA of HFFs infected with parasites harboring a reporter gene (TGME49_243940) expressing GRA81, a merozoite protein endogenously tagged with mCherry within the RH AP2XII-1-mAID- HA or AP2XI-2-mAID- HA lineages. Untreated (UT) and IAA-treated zoites were probed with antibodies against GRA80 (green) and mCherry (red). Cells were co-stained with Hoechst DNA-specific dye. f, M-pileup representation of aligned Nanopore reads at genes identified as up- or down-regulated in clusters A, B, C, or D after IAA-dependent knockdown of AP2XII-1 and AP2XI-2 individually or together.

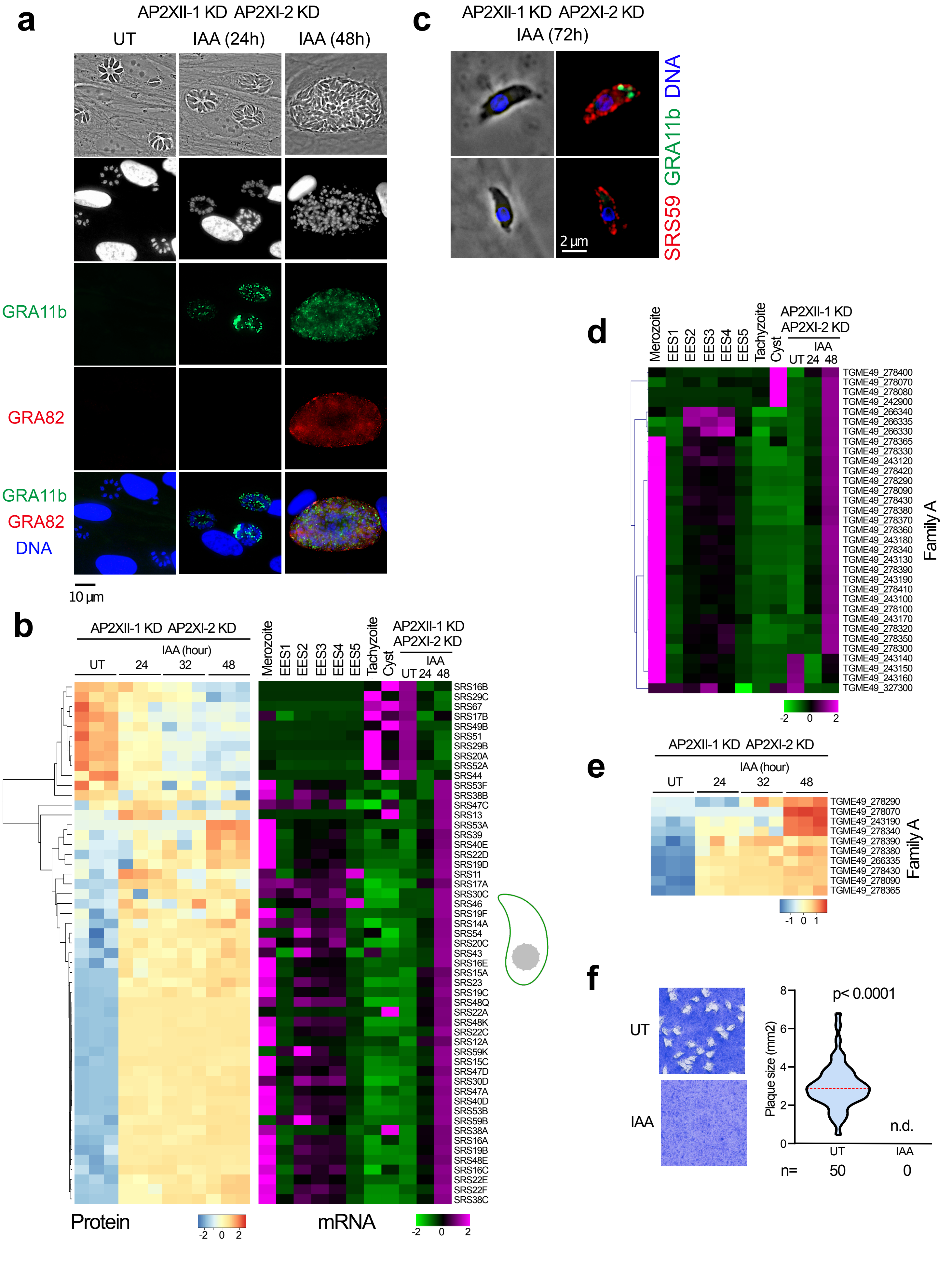


**Extended data Fig. 2**

**Extended Data Fig. 2 | Simultaneous depletion of AP2XI-2 and AP2XII-1 promotes expression of a merozoite repertoire of surface proteins. a**, IFA of HFFs infected with RH AP2XII-1-mAID-HA/AP2XI-2-mAID-MYC parasites. Untreated (UT) and IAA-treated (24 and 48 hours) zoites were probed with antibodies against GRA11b (green) and GRA82 (red). Cells were co-stained with Hoechst DNA-specific dye. **b**, Heat map showing hierarchical clustering analysis (Pearson correlation) of selected SRS mRNA transcripts and their corresponding proteins differentially regulated after simultaneous and conditional depletion of AP2XII-1 and AP2XI-2. Shown is the abundance of their transcripts at different developmental stages, namely tachyzoite, cyst, merozoite, and EES. The color scale indicates the log2-transformed fold changes. **c**, IFA of egressed RH_AP2XII-1-mAID-HA/AP2XI-2-mAID-MYC parasites treated for 72 hours with IAA and probed with antibodies against SRS59 (red) and GRA11b (green). Cells were co-stained with Hoechst DNA-specific dye. **d-e**, Heat map showing hierarchical clustering analysis (Pearson correlation) of selected Family A mRNA transcripts (**d**) and their corresponding proteins (**e**) differentially regulated after simultaneous and conditional depletion of AP2XII-1 and AP2XI-2. Shown is the abundance of their transcripts at different developmental stages, namely tachyzoite, cyst, merozoite, and EES. The color scale indicates the log2-transformed fold changes. **f**, Graphs representing the infectivity of the double KD strain. RH_AP2XII-1-mAID- HA /AP2XI-2-mAID- MYC was either untreated (UT) or treated with IAA for 7 days, and the size of 50 plaques was measured upon detection. Significance was assessed according to Mann-Whitney. n.d., not detected.

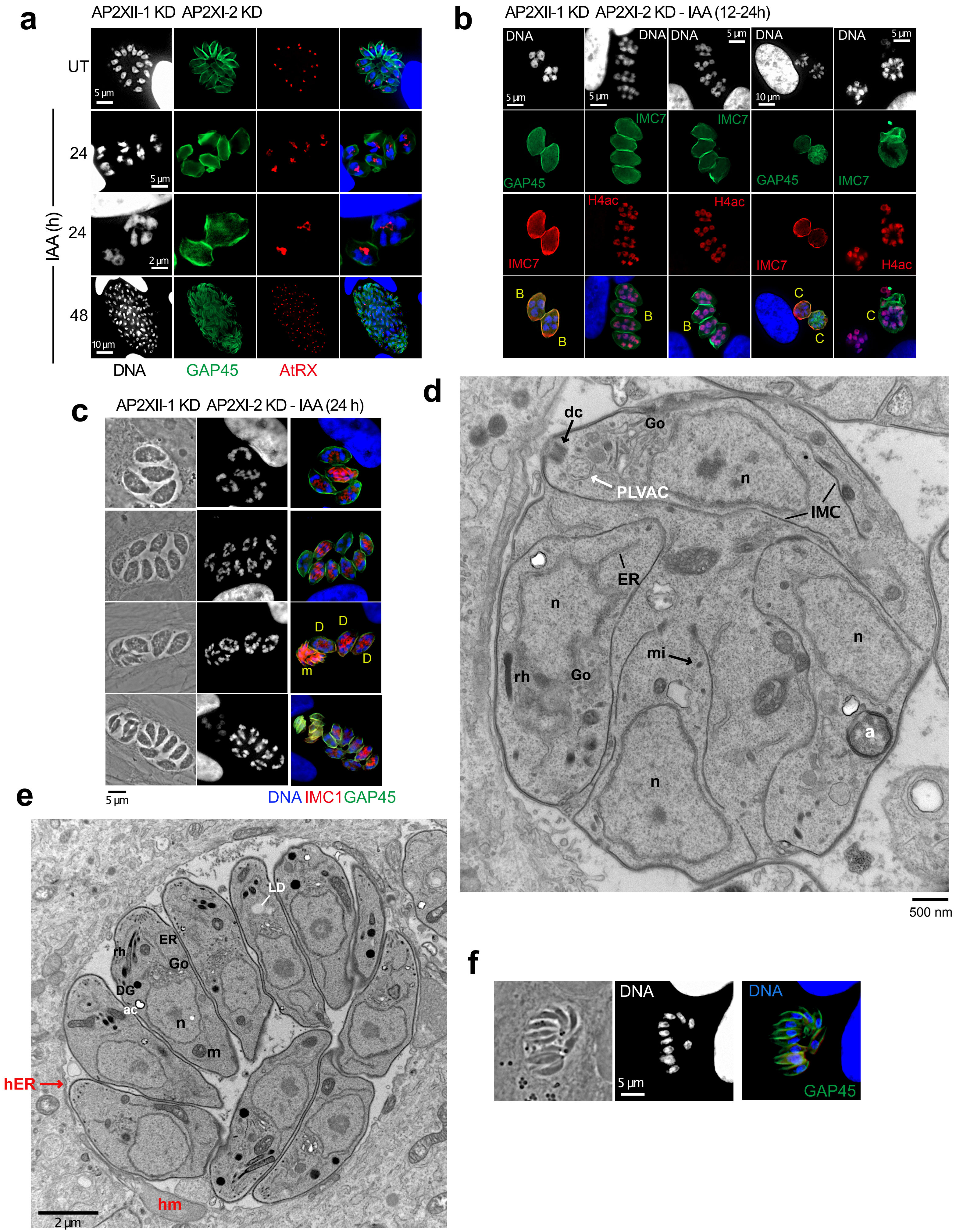


Extended data Fig. 3

**Extended Data Fig. 3 | In vitro development of pre-gametes stages. a-c**, IFA of tachyzoites (UT) and AP2XII-1/AP2XI-2-depleted zoites (at indicated time post-IAA) were fixed and stained with (**a**) antibodies of AtRX clone 11G8 (red) and IMC7 (green), (**b**) IMC7 or pan-acetylated histone H4 (red) and GAP45 or IMC7 (green), (**c**) IMC1 (red) and GAP45 (green). The cells were co-stained with DNA-specific Hoechst dye (white or blue). Types B, C and D meronts are marked in yellow. (**d, e**) Electron micrograph images of RH (AP2XII-1 KD/AP2XI-2 KD)-infected HFFs treated for 24 hours with IAA. **d**, Emphasis on a stage more advance in daughter individualization with the help of the IMC showing polarization with apical conoid. Go: Golgi apparatus, rh: rhoptry, ER: endoplasmic reticulum, mi: microneme, n: nucleus, PLVAC: plant-like vacuolar compartment, dc: daughter conoid, IMC: inner membrane complex. **e**, Emphasis on fully formed merozoites aligned in the PV with apex towards the PV membrane. n: nucleus (plus posterior that in tachyzoite). Go: Golgi apparatus, rh: rhoptry, m: mitochondrion, mi: microneme, ER: endoplasmic reticulum, Go: Golgi apparatus, DG: dense granule, Ac: acidocalcisome, LD: lipid droplet, hER: host endoplasmic reticulum, hm: host mitochondrion. **f**, Fully formed merozoites (24 hours post-IAA) were stained with GAP45 (green) and DNA-specific Hoechst dye (white or blue).

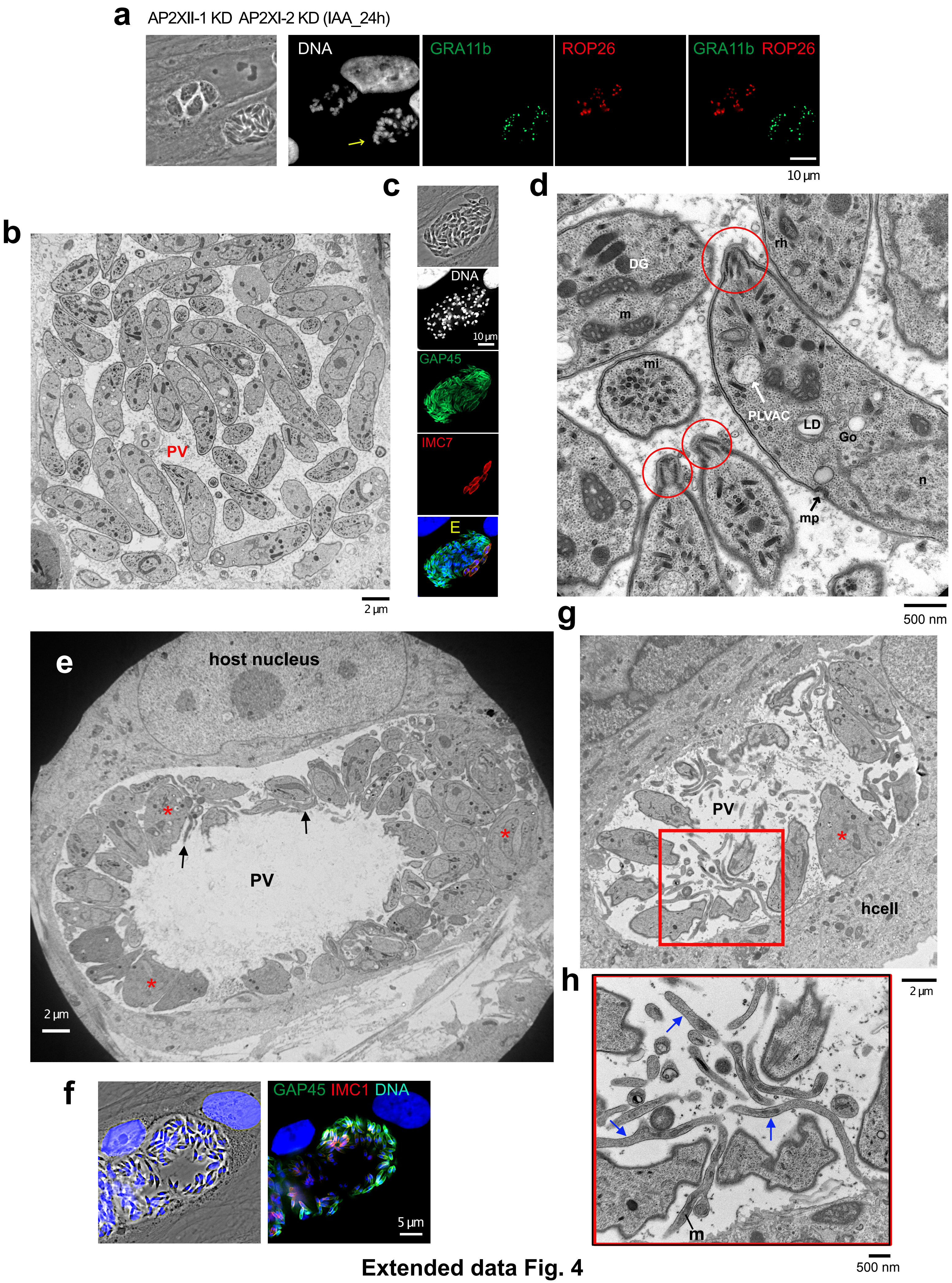


Extended data Fig. 4

**Extended Data Fig. 4 | Mature merozoites and their special substructural features. a**, AP2XII-1/AP2XI-2-depleted meronts (24 hours post-IAA) were fixed and stained with ROP26 (red), GRA11b (green) and Hoechst DNA-specific dye (white). Yellow arrows indicate fully developed merozoites. (**b, d, e, g, h**) Electron micrograph images of RH (AP2XII-1 KD/AP2XI-2 KD)-infected HFFs treated for 48 hours with IAA. b, Emphasis on changes in body shape of merozoite (sausage). **c**, AP2XII-1/AP2XI-2-depleted type E meronts (48 hours post-IAA) were fixed and stained with IMC7 (red), GAP45 (green) and Hoechst DNA-specific dye (white or blue). **d**, Emphasis on conoid extrusion and same organelle content as in tachyzoite. n: nucleus, Go: Golgi apparatus, rh: rhoptry, m: mitochondrion, mi: microneme, DG: dense granule, LD: lipid droplet, mp: micropore, PLVAC: plant-like vacuolar compartment. Red circles showing extruded conoid. **e**, Emphasis on two other morphological transformations of merozoites, either very large (asterisks) or tubular (arrows) in PV with large lumen. **f**, AP2XII-1/AP2XI-2-depleted type E meronts (48 hours post-IAA) were fixed and stained with IMC1 (red), GAP45 (green) and Hoechst DNA-specific dye (blue). (g and h) Emphasis on thin parasitic forms (arrows) containing mitochondria and ribosomes. m: mitochondrion.

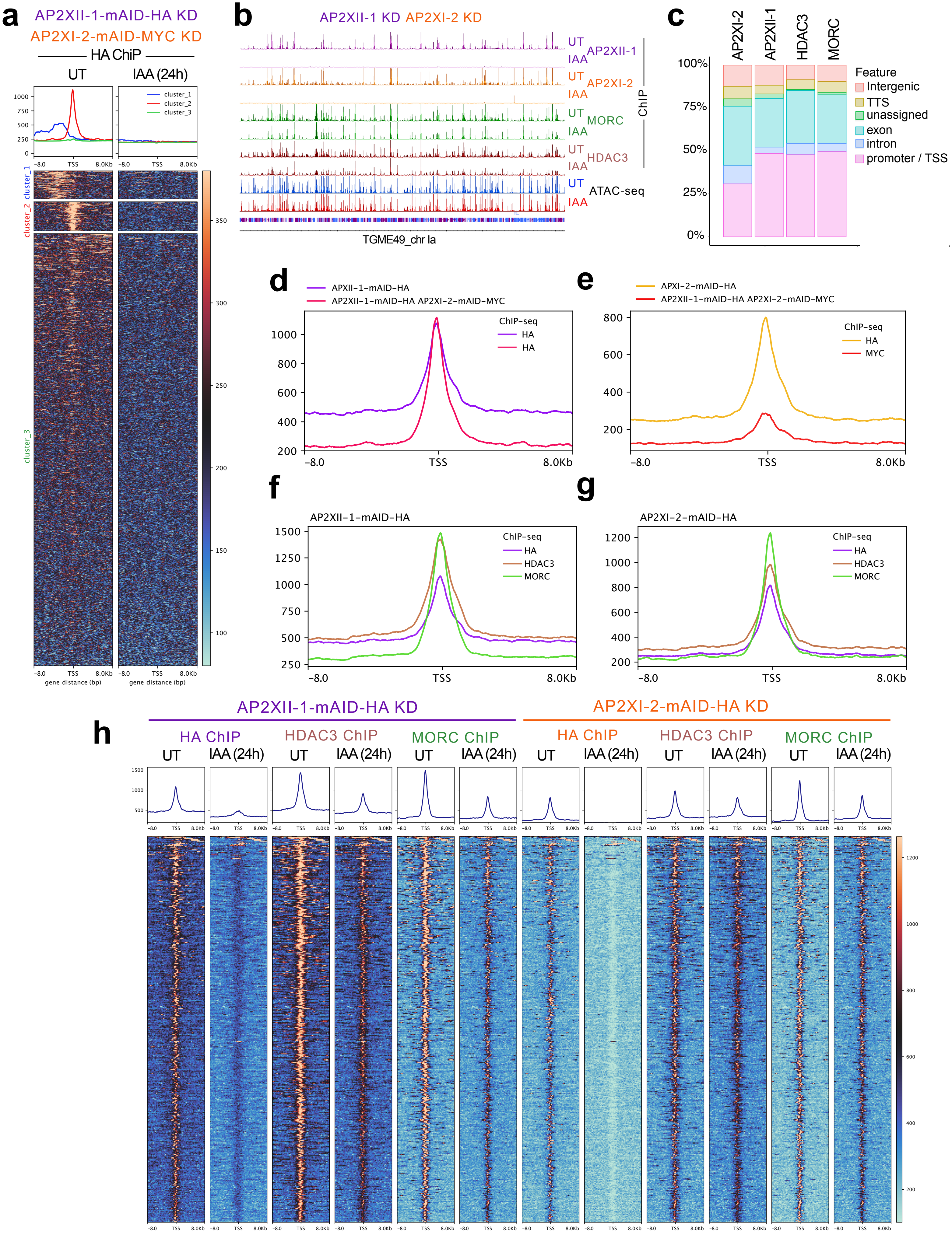


**Extended data Fig. 5**

**Extended Data Fig. 5 | AP2XII-1 and AP2XI-2 jointly recruit MORC and HDAC3 to directly repress the expression of merozoite-specific genes. a**, Heatmap and profile analysis of the chip peak intensity of AP2XII-1 untreated (UT) or after simultaneous knock down of mAID fused AP2XII-1 and AP2XI-2 (IAA 24h). In this case a kmeans clustering is applied on the untreated sample, cluster 2 displaying the most intense and centered genes with AP2XII-1 enrichment at the TSS. **b**, Integrated genome browser view of AP2XII-1, AP2XI-2, MORC, and HDAC3 enrichment on T. gondii chromosome Ia before and after simultaneous knockdown of AP2XII-1 and AP2XI-2. Read density is shown on the y-axis. ATAC-seq chromatin accessibility profiles are plotted for both conditions. **c**, Global distribution of significant chip peaks within genomic features as assessed by HOMER (annotatePeaks) on AP2XI-2, AP2XII-1, HDAC3 and MORC chip experiments. **d**, Comparison profile on cluster 2 (panel A) genomic loci centered at TSS (±8 kb) of chip peaks of AP2XII-1 (using an anti HA chip) within the single knockdown strain of AP2XII-1 or the double knockdown strain of AP2XII-1/AP2XI-2 without auxin treatment. **e**, Comparison profile on cluster 2 (panel A) genomic loci centered at TSS (±8 kb) of chip peaks of AP2XI-2 (using an anti HA or anti MYC chip) within the single knockdown strain of AP2XII-1 or the double knockdown strain of AP2XII-1/AP2XI-2 without auxin treatment. **f**, Comparison profile on cluster 2 (panel A) genomic loci centered at TSS (±8 kb)of chip peaks of AP2XII-1 (HA antibody), HDAC3 and MORC within the single knockdown strain of AP2XII-1. **g**, Comparison profile on cluster 2 (panel A) genomic loci centered at TSS (±8 kb) of chip peaks of AP2XI-2 (HA antibody), HDAC3 and MORC within the single knockdown strain of AP2XI-2. **h**, Profile and heat maps of averaged sum ChIP-seq called peaks showing binding of APXII-1 (HA) or AP2XI-2 (HA), HDAC3, and MORC in the vicinity of TSS of annotated gene promoters before and after addition of IAA for 24h within single knockdown strains of APXII-1 or AP2XI-2. The top panels show the average sum profile on genomic loci centered at TSS (±8 kb). The lower panels show heat maps of peak density around the same genomic loci. The color scale for interpreting signal intensity is on the right side of each graph.

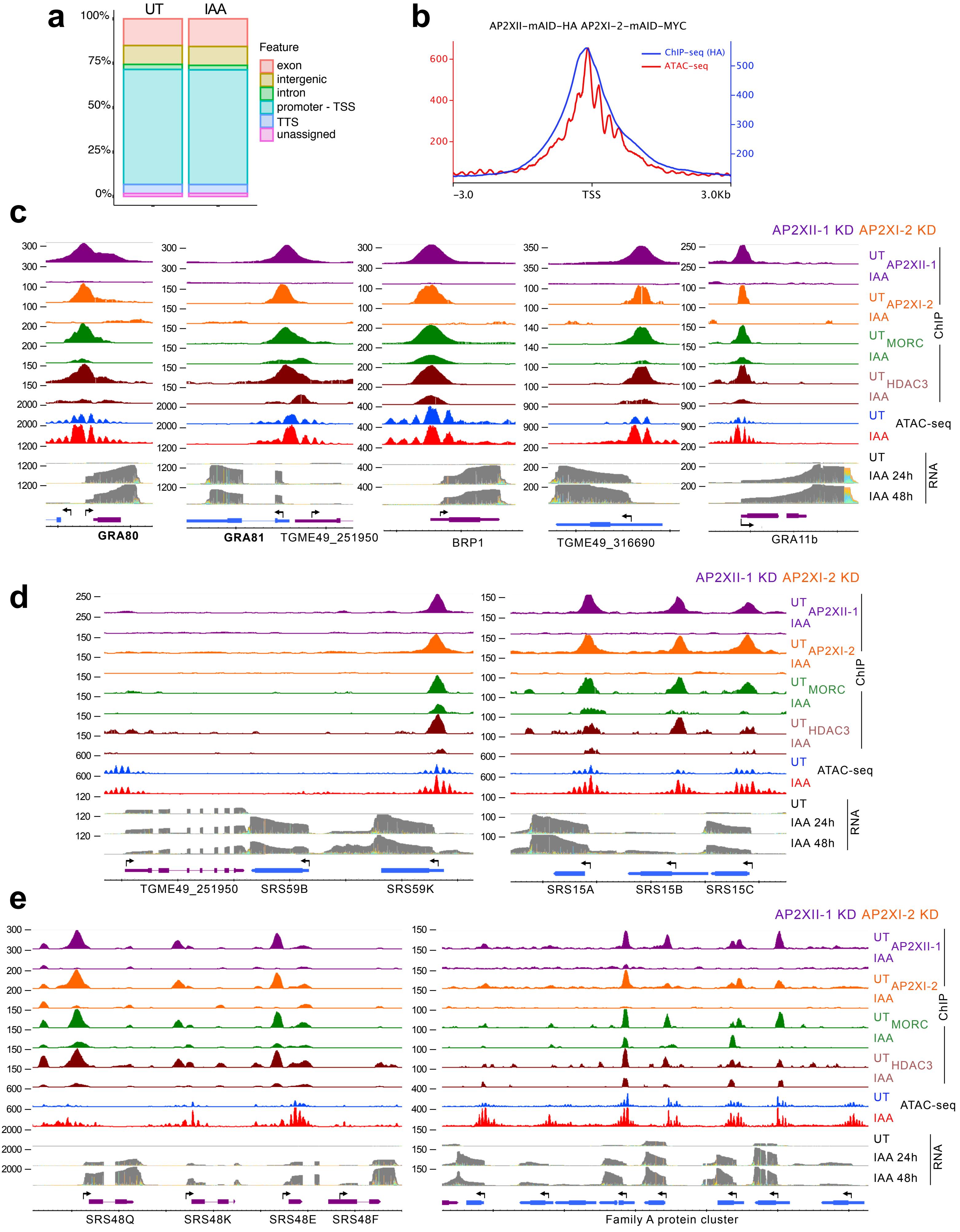


**Extended data Fig. 6**

**Extended Data Fig. 6 | Representative merozoite genes and their regulation by AP2XII-1 and AP2XI-2 and their repressive partners MORC and HDAC3. a**, Global distribution of significant Tn5 accessibility peaks within all genomic features as assessed by HOMER (annotatePeaks) within the AP2XII-1/AP2XI-1 double knockdown strain in untreated or auxin treated conditions. **b**, Comparison profile all genes genomic loci centered at TSS (±3 kb) of chip peaks of AP2XI-2 (HA-antibody) or Tn5 accessibility density within the double knockdown strain of AP2XII-1/AP2XI-2 without auxin treatment. (**c-e**), IGB screenshots of genomic regions with representative merozoite genes. ChIP-seq signal occupancy, ATAC-seq chromatin accessibility profiles, and nanopore DRS are shown in the same manner as in Fig. 6f.

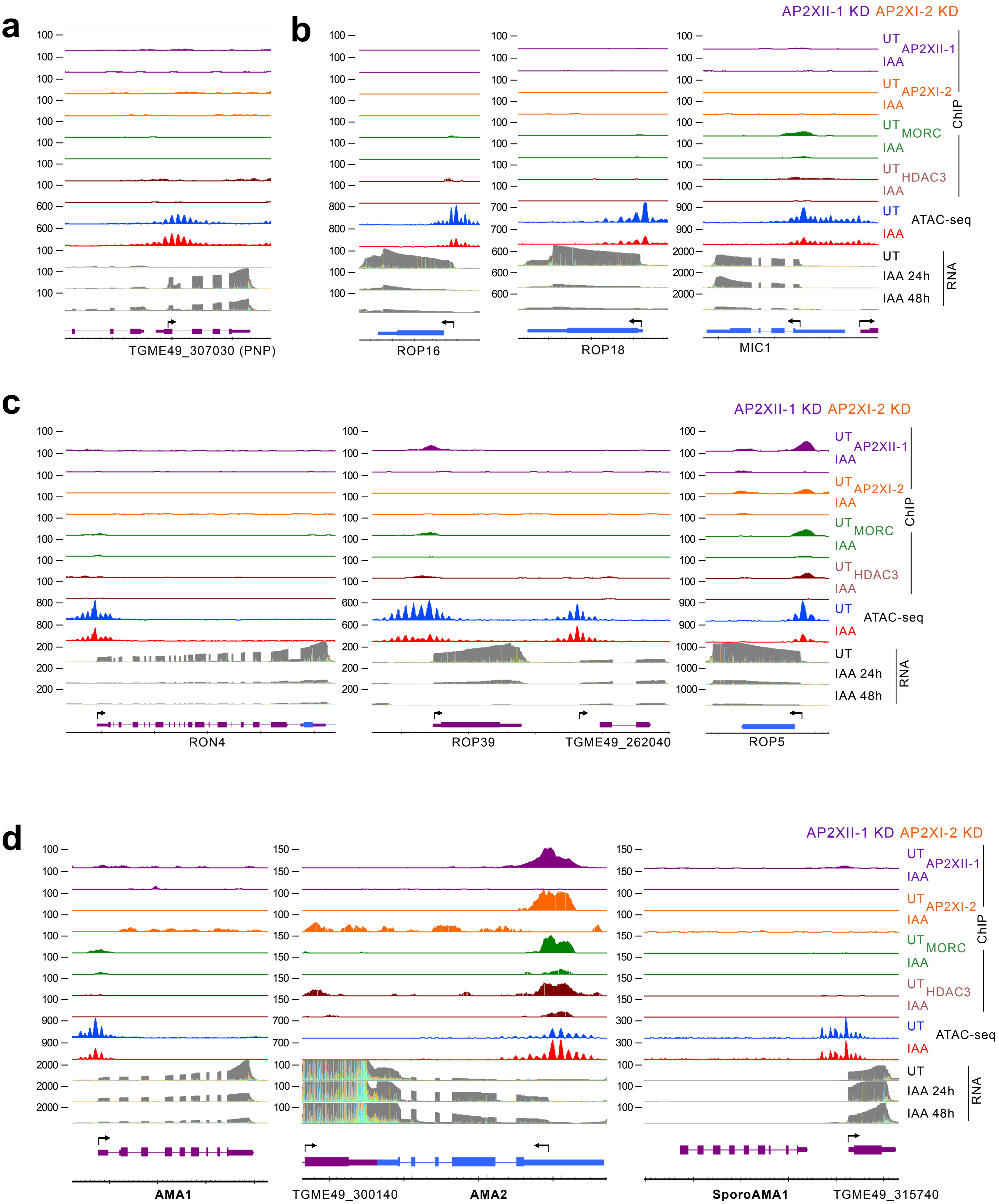


**Extended data Fig. 7**

**Extended Data Fig. 7 | Co-depletion of AP2XII-1 and AP2XI-2 indirectly silences tachyzoite genes**. **a**, IGB screenshot of the genomic region of the merozoite-specific purine nucleoside phosphorylase (PNP) gene. ChIP-seq signal occupancy, ATAC-seq chromatin accessibility profiles, and nanopore DRS are shown in the same manner as in Fig. 6f. **b-c**, IGB screenshots of genomic regions of rhoptry (ROP5, ROP16, ROP18, ROP39, RON4), and microneme (MIC1) genes, shown to be repressed in IAA-treated parasites. ChIP-seq signal occupancy, ATAC-seq chromatin accessibility profiles, and nanopore DRS are shown in the same manner as in Fig. 6f. **d**, IGB screenshots for the AMA1 family are shown with the same caption as in Fig. 6f.

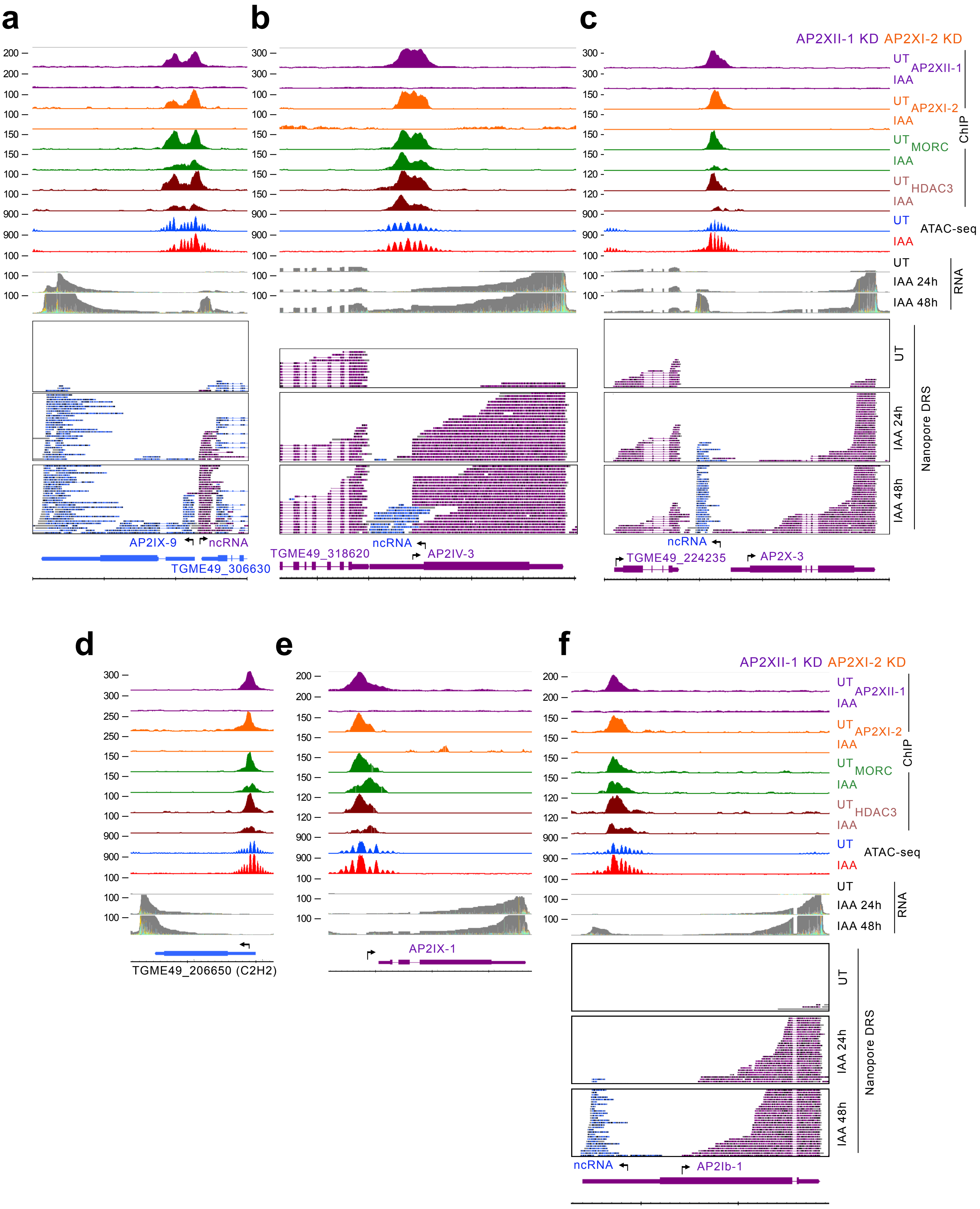


**Extended data Fig. 8**

**Extended Data Fig. 8 | AP2XII-1 and AP2XI-2 co-depletion induces a downstream network of secondary transcription factors to guide merogony. a-f**, IGB screenshots of genomic regions of secondary transcription factors whose expression is activated in IAA-treated parasites. ChIP-seq signal occupancy, ATAC-seq chromatin accessibility profiles, and nanopore DRS are shown in the same manner as in Fig. 6f.
